# Structural influences on synaptic plasticity: the role of presynaptic connectivity in the emergence of E/I co-tuning

**DOI:** 10.1101/2023.02.27.530253

**Authors:** Emmanouil Giannakakis, Oleg Vinogradov, Victor Buendía, Anna Levina

## Abstract

Cortical neurons are versatile and efficient coding units that develop strong preferences for specific stimulus characteristics. The sharpness of tuning and coding efficiency is hypothesized to be controlled by delicately balanced excitation and inhibition. These observations suggest a need for detailed co-tuning of excitatory and inhibitory populations. Theoretical studies have demonstrated that a combination of plasticity rules can lead to the emergence of excitation/inhibition (E/I) cotuning in neurons driven by independent, low-noise signals. However, cortical signals are typically noisy and originate from highly recurrent networks, generating correlations in the inputs. This raises questions about the ability of plasticity mechanisms to self-organize co-tuned connectivity in neurons receiving noisy, correlated inputs. Here, we study the emergence of input selectivity and weight co-tuning in a neuron receiving input from a recurrent network via plastic feedforward connections. We demonstrate that while strong noise levels destroy the emergence of co-tuning in the readout neuron, introducing specific structures in the non-plastic pre-synaptic connectivity can re-establish it by generating a favourable correlation structure in the population activity. We further show that structured recurrent connectivity can impact the statistics in fully plastic recurrent networks, driving the formation of co-tuning in neurons that do not receive direct input from other areas. Our findings indicate that the network dynamics created by simple, biologically plausible structural connectivity patterns can enhance the ability of synaptic plasticity to learn input-output relationships in higher brain areas.

## I. INTRODUCTION

Stimulus selectivity, the ability of neurons to respond differently to distinct stimuli, is one of the primary mechanisms for encoding information in the nervous system. This selectivity can range from simple orientation selectivity in lower sensory areas [1–3] to more complex spatiotemporal pattern selectivity in higher areas [4–6]. Such selectivity is shown to be self-organized under the influence of structured input, enabling, for example, the emergence of visual orientation preference in non-visual sensory areas upon rewiring [7] or changing the whiskers representation in the barrel cortex of rats depending on the level of sensory input [8]. The mechanisms underlying the emergence of input selectivity have been the subject of extensive investigation, both through experimental and computational modelling studies, and still remain under active discussion [9–13].

Despite initially attributing stimulus-selectivity to excitatory neurons and their network structure, we now know that inhibitory neurons are also tuned to stimuli, and the coordination of the E/I currents is a central component of efficient neural computation [14, 15]. In particular, it has been shown that excitatory and inhibitory inputs are often correlated [16], with preferred stimuli eliciting stronger excitatory and, with the small delay, stronger inhibitory responses compared to the non-preferred stimuli [17, 18]. This co-tuning of excitation and inhibition is theorized to be beneficial for a variety of computations such as gain control [19, 20], visual surround suppression [21, 22], novelty detection [23] and optimal spike-timing [15, 24].

Although it is still unclear how E/I co-tuning emerges, the dominant view is that it arises via the interaction of several synaptic plasticity mechanisms [25], a hypothesis that has been reinforced by the findings of multiple theoretical studies over the last decade. First, it has been demonstrated that different inhibitory plasticity rules can match static excitatory connectivity [26–29]. More recently, it was also shown that various combinations of plasticity and diverse normalisation mechanisms allow for the simultaneous development of matching excitatory and inhibitory connectivity in feedforward settings [12, 30–32]. Moreover, a variety of plasticity mechanisms has been associated with the formation of stable assemblies[33–37], the creation of E/I balance [38] and the emergence of tuning selectivity in recurrent networks.

However, so far, most of the theoretical studies of synaptic plasticity have focused on identifying the optimal parameters of individual learning rules and normalization mechanisms for specific tasks and ignore the ways in which these mechanisms act within complex network structures that may influence their function. Specifically, biological networks are characterized by highly non-trivial connectivity structures that are known to display varied degrees of clustering and neuron type-specific connectivity patterns [40]. Such network structures give rise to distinct neural dynamics; for example, clustering in recurrent network structure introduces correlations in the activity of similarly tuned neurons [41] and complex interaction between subpopulations of neurons [42]. These types of dynamics fundamentally alter the statistics of population activity that most synaptic plasticity mechanisms rely on for modifying synaptic strength.

Here, we investigate how the development of matching E/I input selectivity in a downstream neuron via synaptic plasticity can be driven by the structure of recurrent connectivity in the input network. We combine excitatory and inhibitory plasticity rules [26, 31, 43] in the feedforward connections of spiking network to develop detailed cotuning of excitatory and inhibitory connectivity, and we demonstrate that the ability of these plasticity mechanisms to create co-tuning is significantly reduced in the presence of noise and (non-plastic) random recurrent connectivity between the input neurons. We further show that the effects of recurrence and noise on the population activity that drives the formation of matching E/I feedforward weights on a downstream neuron can be fully ameliorated by the introduction of synapse-type specific assemblies of neurons, characterized by local excitation and relatively homogeneous inhibition, an often-observed pattern of cortical connectivity [44, 45]. Our findings demonstrate that network structure can, by influencing population dynamics, significantly modulate the capacity of synaptic plasticity to generate input selectivity in downstream neurons. This highlights a synergistic interaction between structural connectivity and learning mechanisms that can enhance the computational capabilities of brain networks.

## II. RESULTS

We begin by reproducing previously reported results, validating the emergence of co-tuning in a plastic feedforward network, and introducing measures to capture weight diversity and co-tuning for subsequent analysis. Next, we show that the introduction of strong noise or a random static recurrent connectivity in the presynaptic networks impairs the development of co-tuning by destroying the correlation structure in the activity of the presynaptic population. We then illustrate how specific structures in the static recurrent connectivity can restore the ability of plastic synapses to generate co-tuning. Using analytical results from a reduced linear neural mass model and Bayesian inference for the full network, we identify the optimal connectivity structures in static networks, demonstrating that optimal connectivity is influenced by network sparsity. Finally, we simulate fully plastic networks, confirming that our key observations hold true.

### A. Co-tuning and its self-organization by synaptic plasticity in a low-noise feedforward setting

We simulate a single postsynaptic read-out unit driven by a population of *N* = 1000 neurons. The pre-synaptic population is divided into *M* groups *G*_*i*_, *i* ∈ {1, …, *M*}. Each group is comprised of *n* = *N/M* neurons, of which 80% are excitatory and 20% are inhibitory. These neurons are driven by an identical, group-specific Poisson spike train — a shared external input. Additionally, each neuron receives low-intensity independent external noise [26] that prevents unrealistic total synchrony between the input neurons. This setting, depicted in (Fig. 1a), leads to correlated firing among neurons of the same input group (Fig. 1b), which is a common setting for studying the effect of different plasticity rules [12, 26, 31].

**FIG. 1.**
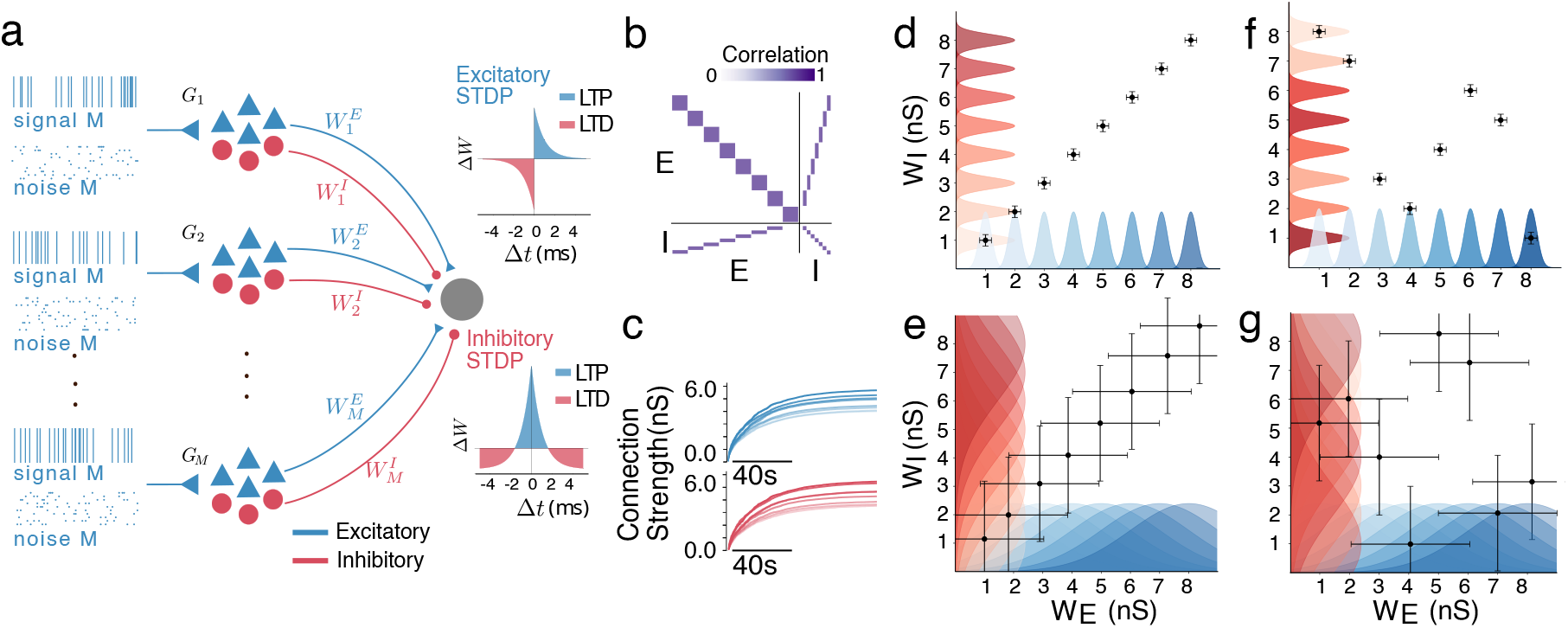
Emergence of co-tuning in a feedforward network. **a**. A diagram of the feedforward network with plastic connections from the different inputs group to the readout neuron.**b**. The correlation matrix of the network’s activity. In the absence of noise, neurons of the same input group are highly correlated. **c**. The development of average E and I weights in an ideal feedforward network with very low noise leads to co-tuned and diverse feedforward connectivity. **d**. An illustration of feedforward connectivity that exhibits both co-tuning and diversity. Different input groups are clearly distinct, and the E and I weights for each group are correlated. The colours of the distribution indicate different groups, and the shading (light to dark) is matched for the corresponding E and I populations; the points and error bars indicate the mean and std of the E and I connectivity of each group. **e**. A co-tuned but not diverse connectivity. E/I weight correlation is maintained, but there is hardly any distinction between groups **f**. A diverse connectivity without E/I weight co-tuning. While each group is distinct from the others, there is no coordination of the E and I connections from the same group **g**. In the absence of weights co-tuning and diversity, the feedforward connectivity lacks any discernible pattern.

Input selectivity develops when the post-synaptic neuron responds differently to inputs from different groups (by adjusting its firing rate). This happens when the average excitatory feedforward projections are sufficiently diverse between pre-synaptic groups (Fig. 1d, f), which leads to groups with stronger feedforward connections eliciting stronger post-synaptic responses upon activation. Moreover, connections from neurons with highly correlated firing (i.e., from the same group) should have a similar strength. To quantify this feature of the network, we define a diversity metric,

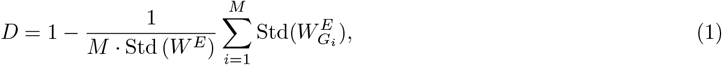

where *W* ^*E*^ ∈ ℝ^*N*^ is the set of excitatory feedforward connection weights, 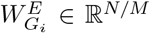 is the subset of excitatory feedforward connection weights from input group *i* and Std(·) denotes the standard deviation. Diversity *D* ∈ [0, 1] equals unity when the feedforward connections from the same group are the same but differ across groups; *D* is close to zero when the feedforward connections from each group follow the same distribution and different groups cannot be meaningfully distinguished (Fig. 1e, g).

Brain networks are characterized by a balance of excitatory and inhibitory currents [18, 46, 47]. For a neuron to be balanced, the average inhibitory current must be equal to the average excitatory current during a (relatively short - usually a few mS) time window. Depending on the temporal resolution of this canceling out, the balance can be more “loose” or “tight”, with detailed (“tight”) balance associated with efficient coding [14, 15] and the ability to encode for multiple stimuli [19] (Further discussion in Supplementary 1). In our specific setting, due to the highly correlated firing in each group, detailed balance can be achieved by matching the relative strengths of the excitatory and inhibitory weights from each group.

To quantify the E/I weight co-tuning, which generates the detailed balance in our simplified network, we use the Pearson correlation coefficient between the mean excitatory and inhibitory weights of each group,

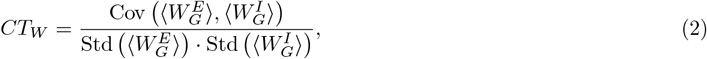

where 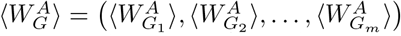, *A* ∈ {*I, E*} and 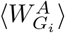 is the average projection weight from the excitatory (*A* = *E*) or inhibitory (*A* = *I*) neurons in group *i*, and Cov(*x*_1_, *x*_2_) denotes the covariance of the variables *x*_1_ and *x*_2_. In networks with strong weight co-tuning *CT*_*W*_, the strength of incoming E and I currents is highly correlated (Fig. 1d, e).

We verify that high diversity (*D* ≈ 1) and weight co-tuning (*CT*_*W*_ ≈ 1) can organically emerge via a combination of plasticity mechanisms in the feedforward connections whose trajectories are initialized at the same (small) value. Specifically, the excitatory connections follow the triplet Hebbian STDP rule [48], and the inhibitory connections follow a symmetric rule [26]. We additionally use competitive normalization in both the inhibitory and excitatory connections, which amplifies small transient differences between the firing rates of different input groups and leads to the development of input selectivity [31]. Since neurons in different groups are independent, while neurons in the same group share a strong correlation (Fig. 1b), the plasticity protocol generates very strongly correlated E/I weights and strong input selectivity, as shown in (Fig. 1c). (For more details on the network’s firing activity, see Supplementary 3).

Postsynaptic weights with high diversity can be well separated, while those with co-tuned weights produce a match between E and I incoming connections. (Fig. 1d-g) illustrates four possible situations. Observe that it is possible to have weights that are, e.g., diverse (so they can be distinguished) but not co-tuned (so E-I are not correlated) and vice versa: E-I weights are correlated, but weights cannot be separated.

### B. Noise and recurrent connectivity compromise the ability of STDP to produce E/I co-tuning

Strictly feed-forward networks with relatively low noise levels are unrealistic approximations of complex cortical circuits (which are characterized by noisy inputs and complex recurrent connectivity), and thus their dynamics might deviate significantly from those observed in experiments. Thus, we introduce noise and non-plastic recurrent connectivity in our pre-synaptic network, both ubiquitously present in biological networks [49, 50]. First, we investigate how they individually affect the emerging E/I co-tuning by changing the structure of the correlations between the neurons of the input network. Then, we examine ways in which different connectivity structures can ameliorate these effects.

We vary the level of noise by changing the fraction of input spikes that are specific to each neuron (noise) vs the shared (signal) input (Fig. 2a). This allows us to control the signal-to-noise ratio while keeping the average number of incoming spikes the same. As the noise intensity increases, the cross-correlations within each input group decrease, while the cross-correlations between neurons of different input groups remain very low, (Fig. 2b). The effect of this in-group decorrelation is an increased variability in the learned projections to the postsynaptic neuron from neurons of the same input group and, thus, a decrease in the resulting diversity (Fig. 2). At the same time, this decorrelation has a much weaker effect on the ability of the plasticity to match E and I feedforward weights from the same groups. This is reflected in the slower reduction of the E/I weight co-tuning, which visibly declines only once the noise becomes overwhelmingly stronger than the input (more than 80% incoming spikes are not shared between neurons of the same group, Fig. 2c). Raster plots illustrating the dynamics for different values of noise are shown in (Fig. 2d).

**FIG. 2.**
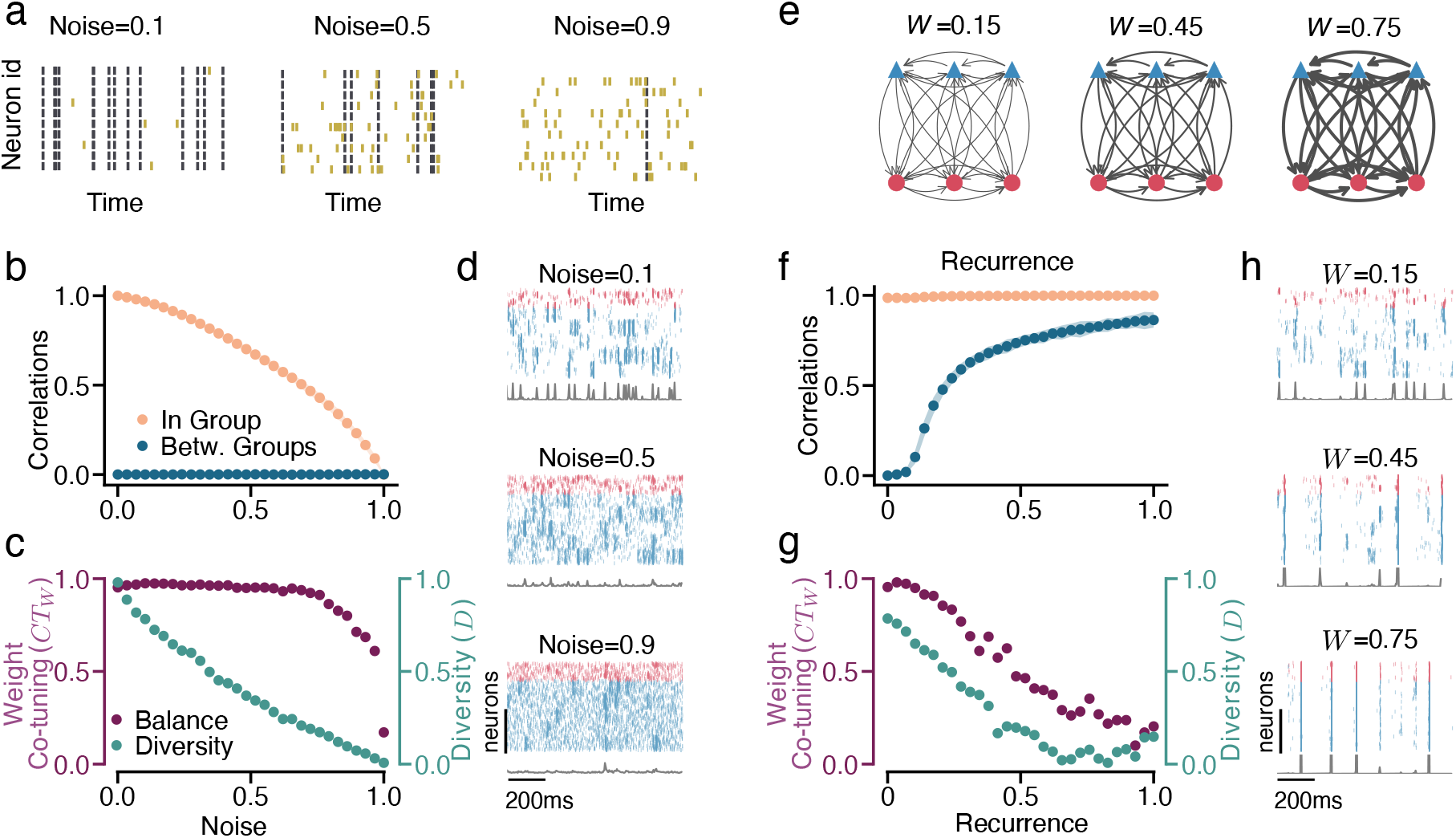
Noise and Recurrence Destroy E/I Co-Tuning. **a**. An illustration of increasing levels of noise in a single input group. In low noise settings, all neurons of the same groups fire at the same time, while as the noise level increases, each neuron fires more and more individual spikes. Joint inputs are shown as black, and individual noise spikes are yellow. **b**. The increase in noise leads to a decrease in in-group correlation (orange), while the between-group correlation (blue) remains low. **c**. An increase in the input noise leads to a reduction in diversity (teal) and, for larger noise intensities, also co-tuning of E/I weights (purple). **d**. Raster plots of input populations’ activity (red, inhibitory neurons; blue, excitatory ones; gray, corresponding firing rate). As the noise increases, the spiking in each group becomes more asynchronous. The traces below (gray line) show the spike count over all neurons in 2ms bins. **e**. An illustration of recurrent connectivity (for three input groups). The coupling parameter *W* controls the mean connection strength. As *W* increases (indicated by thicker connections in the diagram), more of the input a neuron receives comes from other neurons in the recurrently connected network than from the feedforward input. **f**. An increase in the recurrent coupling strength *W* leads to an increase in the between-group correlation (blue), while the in-group correlation (orange) remains high. **g**. The decrease in weight co-tuning and diversity with an increase in the coupling strength. **h**. The spiking activity becomes more synchronous across groups as the coupling strength increases. The traces below (gray line) show the spike count over all neurons in 2ms bins. Simulation parameters not indicated in the text can be found in Supplementary 11

Recurrent connectivity in the pre-synaptic network introduces cross-correlations between the neurons from different input groups, which compromises both diversity and E/I weight co-tuning. To test the extent of this effect, we connect the *N* presynaptic neurons (creating a non-plastic recurrent network) with connection probability *p* and use coupling strength *W* (which denotes the mean synaptic strength) as a control parameter. Initially, we only consider fully-connected networks (*p* = 1). By changing *W*, we can control the ratio between the input received from the feedforward connections (whose rate and connection strength are fixed) and the other neurons in the network via recurrent connectivity (Fig. 2e). The recurrent connectivity increases cross-correlations between groups while maintaining the high correlation within each input group, (Fig. 2f). The effect of these cross-correlations is stronger than the effect of the noise since they affect both the diversity and the weight co-tuning, both of which decline as the recurrent connections become stronger (Fig. 2g). As with noise, raster plots for different recurrent connectivity strengths are shown in (Fig. 2h).

The combination of noise and recurrent connectivity affects both in-group and between-group correlations (Fig. 3c, d), resulting in a reduction of weight co-tuning and diversity as the noise and recurrent connection strength increase, (Fig. 3e, f). The effects of combined noise and unstructured recurrence are not only a sum of their independent effects, but they can lead to novel effects compounding the impact on the emergence of input selectivity. For example, for strong recurrence (Fig. 3d), increasing noise levels seem to lead to an increase of between-group correlations (a counter-intuitive effect, given the decorrelating effects of noise). This is due to the absence of synchronous input to neurons of the same group (due to increased noise, i.e., reduced in-group correlation), which makes synchronisation across groups (and consequently the increase of the between-group correlation) due to recurrent input easier. More details on the effects of noise and recurrence on the network firing are discussed in Supplementary 3 and 4.

**FIG. 3.**
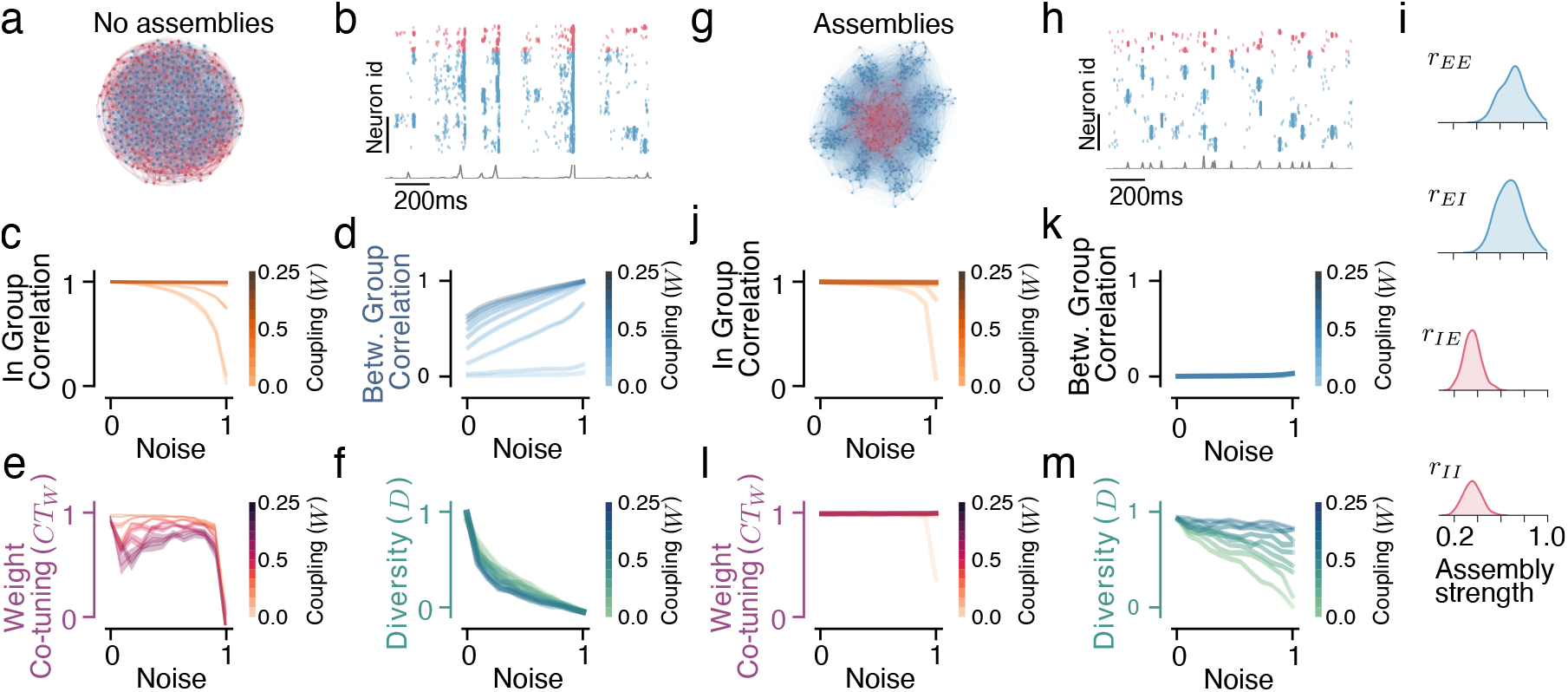
Optimized assemblies of neurons restore the co-tuning in recurrent noisy networks. **a**.Diagram of the network with uniform connectivity **b**. The network activity is characterized by synchronous events across groups. The traces below (gray line) show the spike count over all neurons in 2ms bins. The in-group (**c**.) and between-group (**d**.) correlations for different levels of noise and recurrent connection strength in the uniformly connected network lead to a reduction in the weight co-tuning (**e**.) and weight diversity (**f**.) metrics for different noise and recurrent strength combinations. **g**. Diagram of the network with optimal assembly structure **h**. The network activity becomes more decorrelated across groups. The traces below (gray line) show the spike count over all neurons in 2ms bins. **i**. Approximate posterior distributions of optimal excitatory and inhibitory assemblies strength. The in-group (**j**.) and between-groups (**k**.) correlations for different levels of noise and recurrent connection strength can be almost fully restored by the assembly structure, leading to a restoration of (**l**.) strong weight co-tuning and (**m**.) weight diversity

We develop a formal description of the effect of noise and recurrence on the correlation structure in a simplified linear neural mass model. To this end, we consider *M* = 8 mesoscopic units instead of the previously studied *M* inter-connected groups, represented by continuous rate variables *x*_*j*_(*t*), *j* = 1, …, *M*. These units evolve in time, subject to stochastic white noise. The linear approximation is justified for any system at a stationary state with a constant average firing rate, and it serves as a simplified model for a wide range of parameters of the spiking network (for details on the linear model and its relation to the spiking network, (see Supplementary 6 and 8).

In this simplified case, it is possible to derive analytical equations for all the relevant in- and between-group covariances, which yield the correlation coefficients. These correlations are the solution to a linear system of equations, which can be obtained exactly using numerical methods. Furthermore, one can find close-form solutions in some simple scenarios. For example, in the case of a completely homogeneous network, where all coupling weights are the same, correlation coefficients can be written explicitly (see Supplementary 6 and 8). If the coupling strength increases *W* → + ∞, all correlations grow to 1 as 1 − 𝒪 (1*/W* ^2^). On the contrary, if we reduce the noise *r* → 0_+_ the correlations will decrease to to *MW/*(*M −* 1)^2^ + 𝒪 (*r*^−2^). Both cases eliminate any possible differentiation between the groups, thus compromising the ability of the plasticity mechanisms to create high diversity *D* ≈ 1. Another observation is that in the linear network, increasing noise affects the correlation coefficient quadratically, while coupling increases it linearly. Therefore, since *r <* 1, increasing the coupling has a larger impact on the co-tuning, a consequence that is recovered in the spiking network, consistent with the results shown in (Fig. 2b) and (Fig. 2e).

### C. Neuronal type-specific assemblies restore the ability of STDP to produce co-tuning

The homogeneous all-to-all connectivity (Fig. 3a, b) that we have examined so far is not a realistic assumption and could be particularly detrimental to the self-organization of co-tuning in higher areas. Thus, we examine the impact of different types of inhomogeneous connectivity. In particular, using the idea of functional assemblies (strongly connected neurons that are tuned to the same stimulus [25]), we study whether stronger recurrent connectivity between neurons of the same input group can introduce the necessary correlation structure in the population activity that will allow the plasticity to produce weight diversity and co-tuning.

We maintain the total recurrent input to a neuron constant (the fraction of input coming from the signal/noise vs. the other neurons in the recurrent network, excluding the input from the feedforward connections) while using the ratio of input coming from neurons of the same vs. other input groups as a control parameter. Since we want to vary this ratio independently for each connection type, we define a metric of assembly strength as:

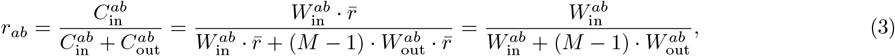

where 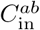 in the total recurrent input a neuron of type *b* receives from neurons of type *a*, for *a, b* ∈ {*E, I*}, of its own input group and 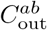 is the total recurrent input the neuron receives from other input groups, 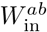 the connection strength between neurons of the same group, 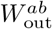 the connection strength between neurons of different groups and 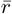 is the average firing rate of network neurons (we assume a uniform firing across the network, which can be simplified out of the equation).

We vary assembly strengths for each type of connection *r*_*EE*_, *r*_*EI*_, *r*_*IE*_, and *r*_*II*_ while keeping the total recurrent input to a neuron 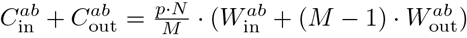 =: *p* · *N* · *W* constant. Here *W* is the average coupling strength, and *p* is the recurrent connection probability. As in the network without assemblies, for now, we consider fully connected networks (*p* = 1). Thus, we can vary the fraction of input coming to a neuron from its own input group without changing the total recurrent E, or I input it receives.

Since the structure of the feedforward connectivity (diversity and weight co-tuning) that the plasticity converges to is controlled by the correlation structure of the inputs, we can use the correlations as a proxy that is easier to optimize than the connectivity metrics. Specifically, we want to maximize the in-group and minimize the between-group correlations, and we seek the assembly structure that leads to that objective. In the reduced linear neural mass model, we compute the optimal assembly strengths (Supplementary 8) analytically and find that strong local excitation and dispersed inhibition restore the desired correlation structure in the network’s activity (Supplementary 9). We find that for all combinations of noise and sufficiently strong recurrent connectivity, strong excitatory assemblies (high *r*_*EE*_ and *r*_*EI*_) and uniform inhibitory connectivity (low *r*_*IE*_ and *r*_*II*_) allow the correlating excitatory currents to remain mostly within the input group/assembly and maintain high in-group correlation, while the diffused inhibitory currents reduce correlations between groups (for a more detailed discussion of this mechanism see Supplementary 10). Still, since the reduced model does not account for many essential features of the spiking network, like sparsity of connections, in-group interactions between neurons of the same type, and non-stationary dynamical states of the groups, the analytic solution obtained for the linear neural mass model can serve to develop intuition, but the results need to be validated for the recurrent spiking network.

We now study the effect of various assembly strengths on weight co-tuning and diversity. Instead of directly assessing it, we turn again to the impact of assembly strength on the spiking network’s activity correlation structure. Thus, we search for combinations *r*_*EE*_, *r*_*EI*_, *r*_*IE*_, *r*_*II*_ that lead to the correlation structure (high in-group and low between-group correlations) that is associated with strong E/I weight co-tuning (*CT*_*W*_ ≈ 1) and maximum weight diversity (*D* ≈ 1). To this end, we use sequential Approximate Bayesian Computation (ABC) [51] to minimize a loss function defined to be zero when the in-group correlations are equal one and all between-group correlations vanish (for details, see Methods).

This method allows us to find the approximate posterior distribution of network parameters (the four assembly strengths) that minimize the loss. Afterward, we verify whether connectivity parameters sampled from the approximate posterior lead to the emergence of diversity and co-tuning in the post-synaptic neuron.

Networks with optimized assemblies largely regain the ability to develop E/I co-tuning despite the noise and the non-plastic recurrent connectivity. Assembly strengths that are drawn from the approximate posterior result in a correlation structure very similar to the one observed in a feedforward/low noise network (Fig. 3j, k), which allows the plasticity to produce a near-optimal structure in the feedforward connections (Fig. 3l, m). We find that the optimal assembly structure involves very strong *E* ← *E* and *I* ← *E* assemblies and medium-strength *E* ← *I* and *I* ← *I* (Fig. 3g, i). For details on the impact of assemblies on the network firing and the learned connectivity, see Supplementary 3 and 4.

This connectivity pattern is similar to the optimal pattern of the reduced linear model, albeit with the difference that the reduced model predicted optimal performance for uniform inhibitory weights (i.e., no inhibitory assemblies, *r*_*IE*_ = *r*_*II*_ = 0). This difference can be attributed to the more complex dynamics of the spiking network that require some degree of local inhibition to prevent extreme synchronization (see Supplementary 7), which can negatively impact the STDP’s ability to produce co-tuning.

This partial specificity of inhibitory recurrent connectivity can be linked to the role of inhibitory tuning in stabilizing network dynamics at the cost of reduced network feature selectivity [52]. In theory, the optimal connectivity pattern in the network would promote competition between different groups, for which completely uniform inhibitory connectivity would be ideal [42]. However, the instability in the population activity induced by such connectivity is detrimental to the emergence of E/I weight co-tuning and input selectivity. Therefore, an intermediate level of specificity in inhibitory recurrent connectivity achieves a balance by maximizing between-group competition (and thus the desired correlation structure) while maintaining stable network dynamics (see Supplementary 7).

### D. The sparsity of a network’s recurrent connectivity shifts the optimal assembly structure

Biological neural networks are usually very sparsely connected [53–55], and the sparsity of connections is associated with distinct dynamics [56]. We observed that the impact of noise and recurrence on the deterioration of weight co-tuning and diversity in sparse networks without assemblies is qualitatively similar to fully connected networks. Thus, we examined the ability of neuronal assemblies to produce activity that restores weight co-tuning and diversity in sparsely connected recurrent networks that receive noisy input.

The optimal assembly strength values depend on sparsity levels. We use ABC to discover the approximate posterior distribution of assembly strengths for 5 different levels of sparsity, corresponding to the probability of connection *p* = 1.0, *p* = 0.75, *p* = 0.5, *p* = 0.25, and *p* = 0.1 (Fig. 4a). We preserve the total input per neuron across different sparsity levels by scaling the coupling strength inversely proportional to *p*.

**FIG. 4.**
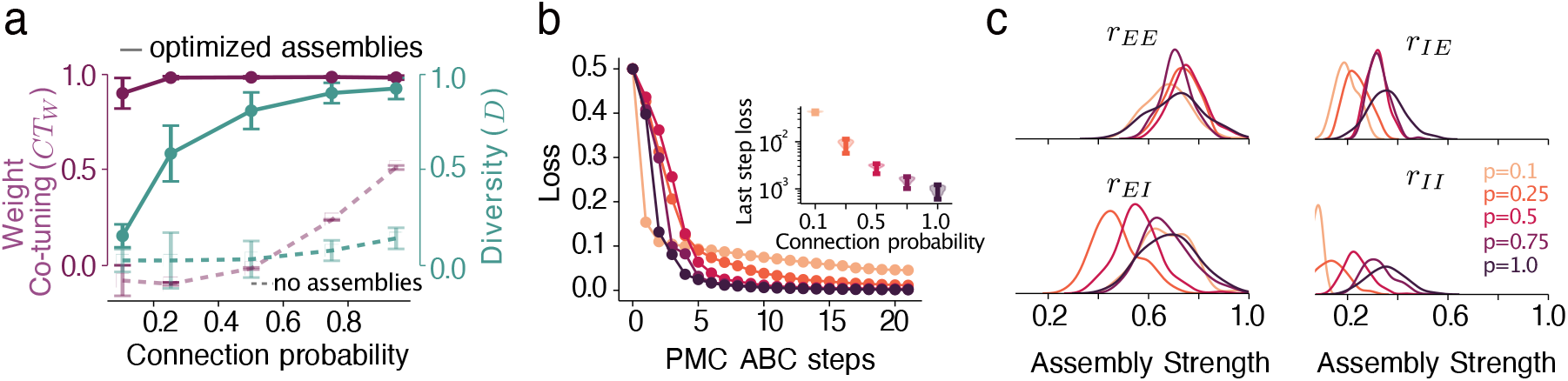
Assemblies improve co-tuning and allow for co-tuning in sparse networks. **a**. weight co-tuning (purple) and diversity (teal) in the networks with assemblies compared to non-structured networks (dashed lines), error bars — standard deviation. The noise level is 0.1 for all sparsities, and the coupling is 1.7 (scaled by 1*/p*). **b**. Loss for the sparser networks is higher, which results in the overall worse performance; the inset shows the loss for 50 accepted samples at the last ABC step. **c**. Posterior distributions of all assembly strengths change with sparsity. Sparser networks require weaker inhibitory assemblies (more uniform connections) to produce co-tuning.

As sparsity increases, the ability of assemblies to improve the tuning diminishes. After 21 ABC steps, the overall loss is larger for the sparse networks than for fully connected networks and increases with sparseness, (Fig. 4b). Therefore, despite an improvement in the tuning metrics for most sparse networks (compare the dashed and solid lines in (Fig. 4a), particularly diversity is strongly affected by sparseness and cannot be recovered by assemblies to the same extent as for the fully connected networks, (Fig. 4a). This reduced effectiveness is expected, given the smaller number of connections and the greater variance in the network’s connectivity.

The optimal strength of most assemblies is reduced as the connection probability is decreased (Fig. 4c). Specifically, we find that all but *E* ← *E* assemblies should be weaker in sparser networks, with the greatest decrease observed in the *I* ← *I* assemblies that completely disappear for very sparse networks. This could be due to a reduced (compared to fully connected networks) need for within-group recurrent inhibition to prevent completely synchronized behaviours.

#### E. Structured connectivity promotes the emergence of co-tuning in fully plastic recurrent networks

In the previous sections, we analyzed a fixed recurrent presynaptic network that projected onto a single postsynaptic neuron via plastic synapses. While this setup provides valuable insights into how structural features can shape population activity for input selectivity to emerge via STDP, it does not fully capture the dynamics of biological neural networks. In this section, we extend our model to a fully plastic and fully recurrent network, offering a more realistic approximation of the behaviour observed in actual biological systems.

In addition to making the recurrent connections between input neurons plastic, we treat the readout neuron as part of the network, and thus, we also introduce sparse feedback connections from the readout neuron to the input neurons (Fig. 5a). Since the recurrent connections are now plastic, we cannot implement the assemblies by changing the in-group connection strength. Thus, we use a sparse network (connection probability *p* = 0.25) and control the in-group connection probability. Following the same logic as with the in-group and between-group connection weights (described in equation 3), we produce in-group and between-group connection probabilities such that the total connection probability (and consequently the total recurrent input) remains constant while the ratio of recurrent input received by a neuron from neurons of its own vs from other groups varies.

**FIG. 5.**
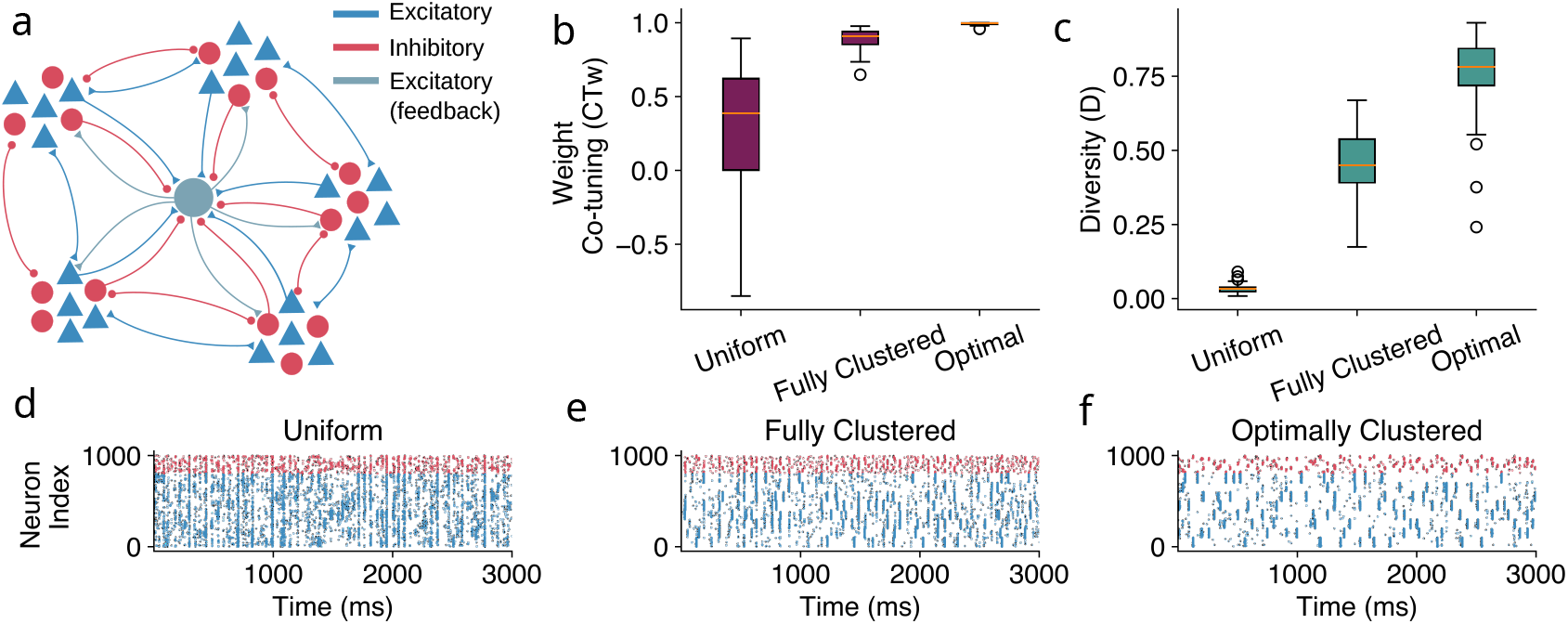
Structured recurrent connectivity can drive synaptic learning even in fully plastic networks. **a**. A diagram of the fully recurrent plastic network. A comparison of the (**b**.) weight co-tuning (*CT*_*w*_) and (**c**.) diversity (*D*) metrics for plastic networks with fully random, fully clustered and optimal connectivity structure. Spike trains of the converged networks with (**d**.) fully random, (**d**.) fully clustered, and (**d**.) optimal recurrent connectivity structures. The activity of different input groups is much more correlated, allowing for easier discrimination in the last network.

We simulate the network with three connectivity structures. A fully random one, where the sparse connections are implemented without any regard for the neurons’ input groups; a fully clustered one, where each input group is almost fully connected, and connections between groups are extremely sparse, and finally, a putative optimally clustered network, which is structured according to the connectivity derived in the previous section for the fixed-weights sparse recurrent network via the Bayesian optimization for the networks of sparsity of *p* = 0.25 (Fig. 4 c). We measure the co-tuning and weight diversity of the readout neuron, which has now been embedded in the network, projecting randomly with plastic connections to the neurons in the presynaptic network with probability *p*.

The fully random network displays very low co-tuning and diversity; the fully clustered network develops stronger co-tuning and diversity, which are then surpassed by the optimally clustered network (Fig. 5b, c). Thus, the putatively optimally clustered network outperforms the other two: after the convergence of the plasticity, the population activity becomes much less noisy, and the activity of different groups can be more easily distinguished, which presumably also enables the development of input selectivity in the post-synaptic neuron (Fig. 5d - f). The correlation structure of the converged plastic networks verifies this observation, with the putative optimally clustered network having very high in-group and near-zero between-group correlation. This structure is introduced at the beginning of the simulation by the structural connectivity and is maintained throughout the learning process, driving the development of input selectivity in the postsynaptic neuron.

Directly optimizing the fully plastic network is not computationally feasible because, for simulation-based inference, we would need to simulate the network’s activity until all the plastic connections converge in each simulation. However, the connectivity from the static sparse presynaptic network appears to give a good prior for a beneficial connectivity in the fully plastic networks.

Our findings suggest that structured connectivity can drive the statistics of activity even in fully plastic networks and enable the development of specific connectivity in neurons that do not receive direct input from other areas.

## III. DISCUSSION

Synaptic plasticity is theorized to be responsible for the formation of input selectivity across brain hierarchies, including in brain areas that only receive input from highly recurrent networks. Here, we demonstrated how the structure of non-plastic presynaptic recurrent connectivity could hinder or boost the ability of synaptic plasticity mechanisms [12, 26, 27, 31] to generate input selectivity in neurons of higher areas. We find that strong excitatory connectivity among neurons tuned to the same input, combined with broader inhibition, creates population activity with a beneficial statistical structure that enables the formation of co-tuned projections by plasticity, potentially fostering input selectivity.

How different plasticity mechanisms shape neural connectivity, such as the formation of E/I co-tuning [12, 26, 27, 31] in feedforward networks or neural assemblies in recurrent networks [26, 34–36], has been a topic of extensive theoretical research. Nevertheless, the opposite effect -the ways in which fixed connectivity can shape the effects of synaptic plasticity - has only been studied in very specialized cases [57]. This omission partially obscures the two-way interaction between connectivity and synaptic plasticity in biological neural networks. While synaptic plasticity constantly modifies some aspects of neural connectivity, it acts within the many constraints of network structure that are either constant throughout an organism’s lifetime or change via structural adaptation mechanisms that act on timescales slower than synaptic plasticity [58, 59]. Effectively, this means that synaptic plasticity relies on population activity originating from networks with highly intricate connectivity structures very different from those of random networks [33, 60]. Given that most synaptic plasticity mechanisms fundamentally depend on the statistics imposed by network activity, it is reasonable to assume that the network structure highly impacts the behaviour of synaptic plasticity.

Cortical connectivity is known to be highly clustered [61], and the clustering has functional as well as spatial determinants. For example, neurons that share common inputs [62] or targets [63] are more likely to form recurrent connections between themselves [64, 65]. Moreover, excitatory cells with similar receptive fields are known to form strong reciprocal connections [66], which determine neural responses. Additionally, cortical networks have been shown to present specific correlation structures early in development [67], suggesting that recurrent cortical connectivity is at least partially structured before sensory inputs are present.

Clustered networks display distinct dynamics, including competition between clusters [42, 68] and slower timescales[41, 69], both of which can be useful for computations. Additionally, there is strong evidence that groups of highly interconnected neurons (neuronal assemblies) share common functions within recurrent networks [37, 70, 71]. Moreover, evidence has accumulated [72, 73] that different neuron types (excitatory and inhibitory subtypes) follow distinct spatial connectivity patterns, which have implications for neural computation [74]. Thus, our findings complement the ongoing research on computational implications of recurrent neural connectivity in biological networks by suggesting a link between specific, fixed connectivity patterns and local learning in feedforward projections to downstream populations via synaptic plasticity.

While the connectivity pattern we identify in our study is biologically plausible, the extent to which it is realized *in vivo* remains unclear. Further experimental studies in the connectivity patterns of different neuron types are necessary to model different network connectivities and study their dynamics effectively. On a theoretical side, more research is needed to uncover whether the synapse-type specific assembly structures we identify can emerge in plastic networks without any prior structure. Specifically, while there have been several studies investigating the emergence of stable assemblies via STDP [33–35, 75], the protocols by which assemblies of different pre-selected strengths for each connection type could arise remain unclear. Potential mechanisms include structural plasticity and variation in the learning rates of different synaptic types. Additionally, the presence of different regulatory interneurons, which have already been studied in the context of assembly formation [34], could play a role in modulating the relative assembly strengths of different connections.

For our study, we parameterized the network connectivity by adopting a quantitative metric for the strength of different types of neuronal assemblies. This resulted in a low-dimensional parameter space and allowed us to use rejection sampling-based ABC [51] to infer the optimal assembly strengths. One limitation of this technique is that it suffers from the curse of dimensionality and typically requires a large simulation budget [76, 77]. Therefore, extensions of the current work using higher dimensional connectivity parameters or simultaneous inference of the connectivity and neuron parameters will require more efficient simulation-based methods such as neural posterior estimation [78]. Alternatively, direct optimization of each recurrent weight via gradient-based methods [79, 80] may uncover more intricate connectivity patterns that are not limited to the specific network parametrization we chose.

To summarize, we identified how particular presynaptic connectivity structures could be a favourable or detrimental substrate for plasticity to develop co-tuning of excitation and inhibition on neuronal projections. Our study is the first step in illuminating the two-way dependence between the non-plastic structural features of a network’s connectivity and synaptic plasticity, which can motivate further research on this intricate interaction.

## IV. MATERIALS AND METHODS

### A. Neuron model

We modelled all neurons of our networks as leaky integrate-and-fire (LIF) neurons with leaky synapses[81]. The evolution of their membrane potential is given by the ODE:

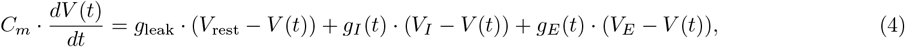

where *V*_rest_ is the neuron’s resting potential, *V*_*E*_, *V*_*I*_ are the excitatory and inhibitory reversal potentials and *g*_leak_ the leak conductance. Additionally, the excitatory and inhibitory conductances *g*_*E*_, *g*_*I*_ decay exponentially over time and get boosts upon excitatory or inhibitory pre-synaptic spiking, respectively, as

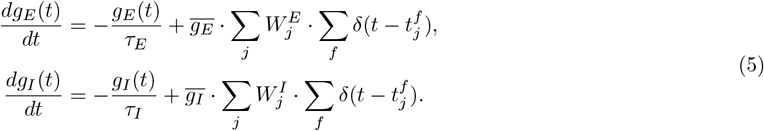

Here 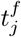 denotes the time at which the *f* -th spike of the *j* − th neuron happened and *δ*(*t*) is the Dirac’s delta function. When the membrane potential reaches the spiking threshold *V*_th_, a spike is emitted, and the potential is changed to a reset potential *V*_reset_. Finally, the neurons have an absolute refractory period (during which no spikes are emitted even when the spiking threshold is reached) between spikes of *τ*_ref_ = 5*ms*.

We would like to remark that both 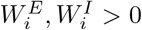, since the effect of inhibition is encoded on eq. (4). However, for illustration purposes inhibitory weights are currents are shown to be negative. For instance, this happens in (Fig. 1b). An alternative neuron model is discussed in Supplementary 2.

### B. Network input

The external input to each of the 1000 pre-synaptic neurons is the mixture of two Poisson spike trains. The first Poisson spike train is shared with all the other neurons of the same group, while the second Poisson spike train is the individual noise of the neuron,

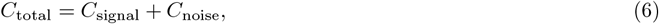

where *C*_signal_ ∼ Poisson((1 − *c*) · *f*_0_) and *C*_noise ∼_ Poisson(*c* · *f*_0_). Here, *f*_0_ is the total firing rate of the input, and *c* is the strength of the noise. *C*_signal_ is the same for all neurons of the same input group, while *C*_noise_ is individual to each neuron

### C. Recurrent connectivity

The recurrent connectivity is implemented in two different versions for the fixed and plastic versions.

#### 1. Non-plastic recurrent connectivity

The non-plastic recurrent connectivity between the input neurons is defined by a coupling strength parameter *W*, which defines the average synaptic strength and a connection probability *p*, which defines the sparsity. The connectivity is implemented as follows:

At first, an adjacency matrix *A* is defined, which implements an Erdős–Rényi connectivity with connection probability *p* (i.e., a connection between any two neurons is implemented with connectivity *p*, independently of any other connection). Then, using the coupling strength parameter *W*, and the given assembly strength for each connection type *r*_*EE*_, *r*_*EI*_, *r*_*IE*_, and *r*_*II*_, we extract the parameters 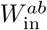 and 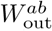 for each connection type (*a, b* ∈ {*E, I*}) according to eq 3. The inhibitory weights 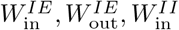 and 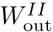 are scaled by a parameter *g*_*s*_ which is set to counterbalance the slower inhibitory synapse dynamics and the smaller number of *I* neurons. This scaling leads to an approximately balanced network across implementations. Once the connectivity strengths are calculated, for each pre and post-synaptic neuron pair *i* and *j*, we set the connection between them as

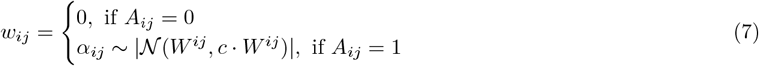

where *W*^*ij*^ is the appropriate connectivity strength (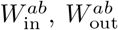 for *a, b* ∈ {*E, I*}) depending on the neuron type of neurons *i* and *j* and whether they belong to same assembly or not. The parameter *c*, which scales the standard deviation, was normally set to 0.1, but we also examined narrower and broader distributions with similar results.

We finally considered an alternative, log-normal distribution of weights, which increased variability but largely lead to the same results.

#### 2. Plastic recurrent connectivity

In the plastic recurrent network case, instead of varying the connection strengths for in-group and between-group connections, we use the total connection probability *p* and given assembly strength for each connection type *r*_*EE*_, *r*_*EI*_, *r*_*IE*_, and *r*_*II*_, we extract parameters 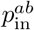and 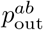 for each connection type (*a, b* ∈ {*E, I*}) according to eq 3. These parameters give the probability that a connection of a specific type is implemented in the recurrent network, which creates an inhomogeneous adjacency matrix *A* that implements the different levels of clustering for each connection type.

Once the adjacency matrix has been defined, we set the initial connection strength for pre and post-synaptic neuron pair *i* and *j* as :

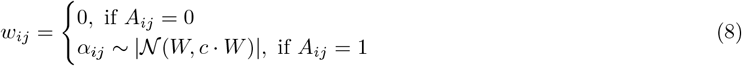

where *W* is the coupling strength parameter, which we scale for inhibitory connections, similarly to the non-plastic network. The resulting connectivity reflects the initial conditions for the plastic recurrent network and the non-zero connections are updated according to the same plasticity protocol that is used to learn the feedforward connectivity in the networks with fixed recurrent connectivity.

### D. Plasticity

#### 1. Triplet excitatory STDP

The excitatory connections are modified according to a simplified form of the triplet STDP rule [43], which has been shown to generalize the Bienenstock–Cooper–Munro (BCM) rule [9] for higher-order correlations [48]. In our implementation of the triplet rule, the firing rates of the pre-synaptic excitatory neurons and the post-synaptic neuron are approximated by traces with two different timescales (we use the same timescales for the fast and slow traces of the pre and postsynaptic neurons):

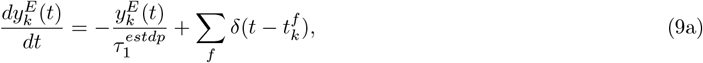

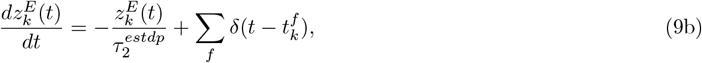

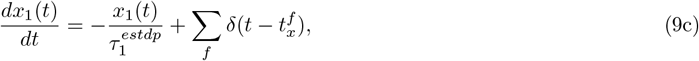

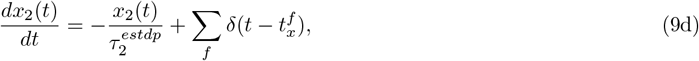

where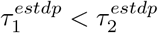 are the two timescales of the plasticity rule, 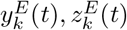 and 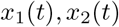 represent the slow and fast traces of the *k*-th excitatory pre-synaptic and the single post-synaptic neuron respectively while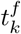and 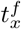 are their respective firing times The function *δ*(*x*) represents a Dirac’s delta. The connection weights are updated upon pre and post-synaptic spiking according to

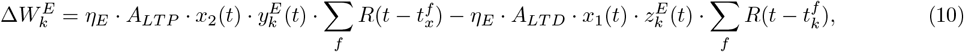

where *η*_*E*_ is the excitatory learning rate and *A*_*LTP*_, *A*_*LTD*_ the amplitudes of long term depression and potentiation respectively. Despite scaling down the LTD amplitude to account for higher presynaptic firing rates, the rule remains slightly LTD dominated in our experiments, a setting that has been observed in experimental studies [82, 83]. The function *R*(*x*) is defined as:

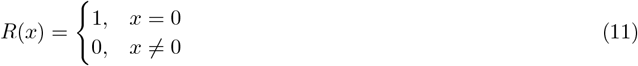

In numerical simulation, if spikes have been produced in the last few timesteps, so in practice *R*(*t*) = 1 for *t* ∈ [− *δt*, 0] for a small *δt*. Hence, *R*(*t*) is a rectangular function. Different parameters and learning rules for the excitatory plasticity are discussed in (Supplementary 5.A and 5.B).

#### 2. Inhibitory STDP

We used the learning rule first proposed in [26] for the inhibitory connections. Approximations of the firing rates are kept via a trace for each of the pre-synaptic inhibitory neurons as well as the post-synaptic neuron,

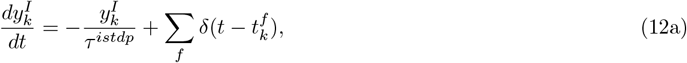

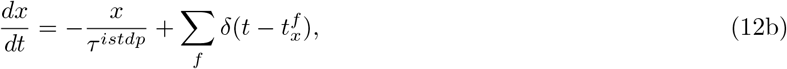

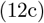

where *τ* ^*istdp*^ is the single timescale of the plasticity rule, 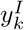and *x* are the traces of the the *k*_*th*_ inhibitory pre-synaptic and the single post-synaptic neuron, and 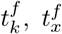 are their respective spike times. The connection weights are updated upon pre and post-synaptic spiking as

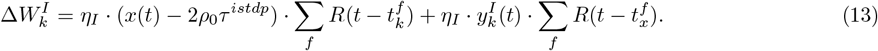

Here, *η*_*I*_ is the inhibitory learning rate, and *ρ*_0_ is the target firing rate of the post-synaptic neuron. The rectangular function *R*(*t*) is defined in eq. (54).

#### 3. Weight normalization

Due to the instability of the triplet STDP rule, some normalization mechanism is needed to constrain weight development. We use a modified version of the competitive normalization protocol proposed in [31], which we adapt for spiking neurons.

Specifically, we normalize the *k*-th connection every time there is a weight update (i.e., upon pre or postsynaptic spiking):

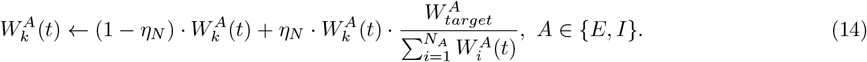

Where 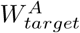 is the target total weight of each connection type and *η*_*N*_ is the normalization rate. In the recurrent plastic networks, the 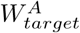 for the recurrent neurons is determined by the coupling strength *W*. The normalization pushes the sum of the excitatory and the sum of the inhibitory feedforward connections weights close to the set target total weights 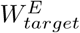 and 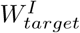 over time. The implications of implementing a regular normalization step (on every time step only when spiking occurs) are discussed in (Supplementary 5.D.1). Moreover, the implications of using different normalization rates *η*_*N*_ are discussed in (Supplementary 5.D.2)

An alternative way to stabilize the weights via subtractive normalization of only the excitatory synapses [12, 23, 33] was also considered leading to comparable results (see Supplementary 5.E).

### E. Approximating the posterior distribution of the model parameters

To estimate the set of parameters that lead to high in-group correlations and low out-group correlations, we used simulation-based inference [76]. The basic idea is to use simulation with known parameters to approximate the full posterior distributions for the model given the required output, i.e., the distribution of parameters and samples from which produce the required correlation structure. We use sequential Approximate Bayesian Computation (ABC) [51] to approximate the posterior distribution. We define a loss function that maximizes in-group correlations and minimizes between-group correlations:

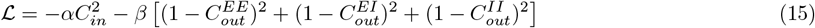

We define a uniform prior *p*(*θ*). A set of parameters *θ* = [*r*_*ee*_, *r*_*ei*_, *r*_*ie*_, *r*_*ii*_] is sampled from it and used to run the simulations for 3 seconds. From the simulation results, correlations are computed, which allows us to obtain the loss. We accept a parameter set if the loss is below the error *ϵ*, and keep sampling until the number of accepted samples is We use the kernel density estimate on the accepted samples to obtain an approximate posterior. Next, we rescale this approximate posterior with the original prior to obtain a proposal distribution that we use as a prior in the next step of the ABC. In each step, we reduce *ϵ* by setting it to the 75th percentile of the losses for the accepted samples (see [51] for more details). As a rule, we run 20 to 30 steps of the sequential ABC until the loss converges. We run separate fits for networks with different levels of sparsity with connection probabilities *p* = 0.1, 0.25, 0.5, 0.75, 1.0. The fitting was done using a modified version of the simple-abc toolbox https://github.com/rcmorehead/simpleabc/ for python.

### F. Reduced model

The dynamics of the system can be studied analytically using a simplified, reduced linear model. Here, each pair of variables (*x*_*i*_, *y*_*i*_) represents the excitatory and inhibitory mean firing rate of a neuron group. In theory, these variables display complicated non-linear interactions that arise from the microscopic details of the LIF spiking network and synapse dynamics. However, in the stationary state –and away from any critical point– a linearised model can capture the essential features of the correlations between different populations.

Internal noise, modelled as independent Poisson trains to each individual neuron, becomes Gaussian white noise in the large-population limit, characterized by zero mean and variance *σ*_int_. Each population is affected by different internal fluctuations. For simplicity, external noise, which is applied as the same train of Poisson spikes to all the neurons inside an input group, will also be approximated as a Gaussian white noise of mean *η*_0_ and variance *σ*_ext_.

Therefore, the simplified linear model reads:

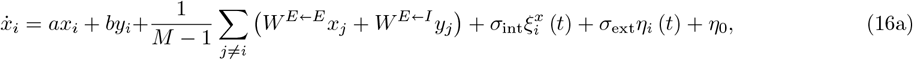

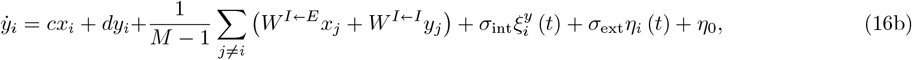

where *M* is the number of populations, *a, b, c, d* are parameters controlling in-group recurrent coupling, and *W* ^*E* ← *E*^, *W* ^*E* ← *I*^, *W* ^*I* ← *E*^, *W* ^*I* ← *I*^ are couplings between different clusters. Internal noise for each population is represented by 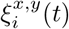, while external noise is notated as *η*_*i*_(*t*). All noises are uncorrelated, meaning that

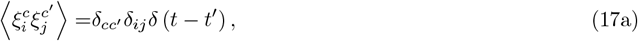

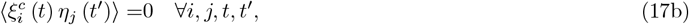

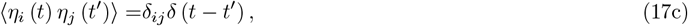

with *c, c*′ = {*x, y*}, and where ⟨…⟩ represents an ensemble average, i.e., an average over noise realizations. From this model, it is possible to obtain closed equations for Pearson correlation coefficients (see Supplementary 8 for details). Notice that stochastic differential equations are never complete without an interpretation, and we choose to interpret these in the Itô sense, which will be relevant for computations. Tables of all the parameters used in our simulations are given in Supplementary 11.

## V. ACKNOWLEDGEMENTS

This work was supported by a Sofja Kovalevskaja Award from the Alexander von Humboldt Foundation. EG thank the International Max Planck Research School for Intelligent Systems (IMPRS-IS) for support. We acknowledge the support from the BMBF through the Tübingen AI Center (FKZ: 01IS18039B). AL is a member of the Machine Learning Cluster of Excellence, EXC number 2064/1 – Project number 39072764.

## Appendix 1 E/I co-tuning and input selectivity

In our study, we use the weight diversity as a proxy for input selectivity. Verify that this mapping is reasonable even for networks with very detailed E/I balance. Specifically, we model 2 networks with fixed, perfectly balanced connectivity. The first of the two has tuned excitation (Fig. 6a) and totally flat inhibition (leading to loose balance), while the second has near-perfect co-tuning of its incoming *E* and *I* incoming weights, which leads to tight balance (Fig. 6b).

**FIG. 6.**
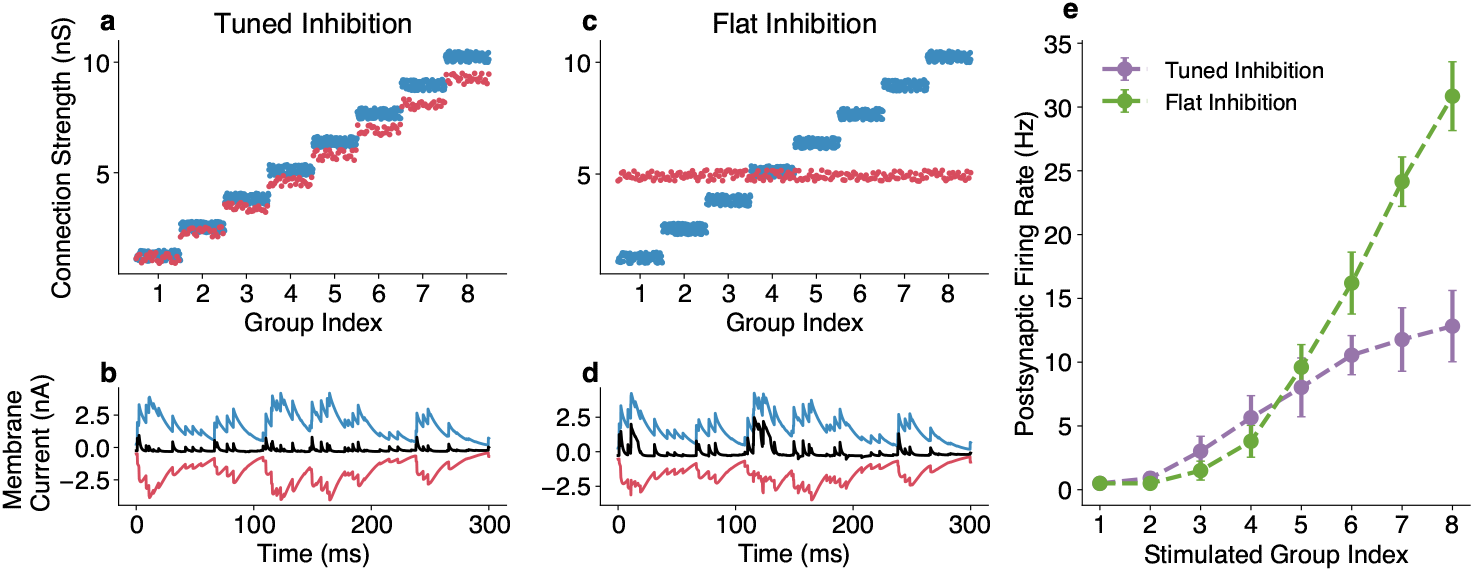
E/I Weights and input selectivity: **a**. The feedforward connectivity and **b**. the resulting postsynaptic currents for a network with flat co-tuned *E/I* feedforward weights feedforward inhibition. **c**. The feedforward connectivity and **d**. the resulting postsynaptic currents for a network with flat feedforward inhibition. **e**. The firing of the postsynaptic neuron upon activation of different input groups for each network. Flat inhibition leads to sharper responses while co-tuning leads to lower FR and a more gradual increase of activity for different input activation.

We sequentially activate different input groups and record the post-synaptic neuron’s response in terms of incoming currents and firing rate. We see that while there is a small difference between the incoming currents in the two cases (Fig. 6b and d), the firing rate of the postsynaptic neuron is significantly different between the two networks (Fig. 6e).

In particular, we see that compared to the network with flat inhibition, the responses of the network with the co-tuned *E/I* weights are significantly reduced but still clearly different for different inputs, allowing discrimination. This is consistent with recent theoretical findings [52] about inhibitory tuning reducing input selectivity.

However, the co-tuned network displays a reduced firing rate, which suggests the possibility for more efficient encoding as well as a partial encoding for non-preferred inputs (i.e., a small but detectable response for the input group associated with small input weights) in contrast to the neuron with flat inhibition, which is fully silent for the non-preferred inputs (i.e., those for which it receives stronger inhibition than excitation). This agrees with earlier findings on the benefits of weights co-tuning in terms of efficiency and coding [14].

This suggests that while the detailed co-tuning of feedforward *E/I* weights might lead to a reduction in the sharpness of the post-synaptic neuron’s tuning, it still enables discrimination between inputs and, indeed, can encode for a broader range of inputs (responses even for non-preferred stimuli) in a more energetically efficient way (lower firing rate).

## Appendix 2 The effects of inhomogeneous connectivity on the population activity are independent of the neuron model

In order to ensure that our results are independent of the exact neuron model we are using, we repeated the bayesian fitting with a simplified network of LIF neurons with current based synapses.

Specifically, the voltage is given by:

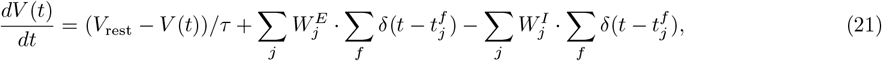

where *V*_rest_ is the neuron’s resting potential and *τ* is the membrane timescale. Here 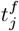 denotes the time at which the *f*-th spike of the *j*–th neuron happened. When the membrane potential reaches the spiking threshold *V*_th_, a spike is emitted, and the potential is changed to a reset potential *V*_reset_.

We repeat the fitting with ABC for a network with this model, using the same objective function (maximizing in-group and minimizing between-group correlations) and we find that the optimal assembly strength distribution for all four connection types we identified for the original network remains very similar to the ones we identified for the original model.

## Appendix 3 Firing behaviour of the presynaptic network

In order to better understand the effects that the introduction of noise and non-plastic recurrent connections has on the population activity of the presynaptic network, we calculate a number of metrics that quantify population activity for different connectivity structures.

At first, we look at a feedforward network with relatively low noise (noise intensity is set to 0.15). This setting produces activity that is ideal for the plasticity to produce diverse and co-tuned E/I weights.

As expected, due to the Poisson input and the absence of any recurrent interactions, the networks have a broad ISI distribution, a relatively narrow distribution of the CV of the ISI and a Fano Factor very close to 1 (Fig. 7a-c). The activity of the network is highly correlated within groups, but different groups fire independently (Fig. 7d).

**FIG. 7.**
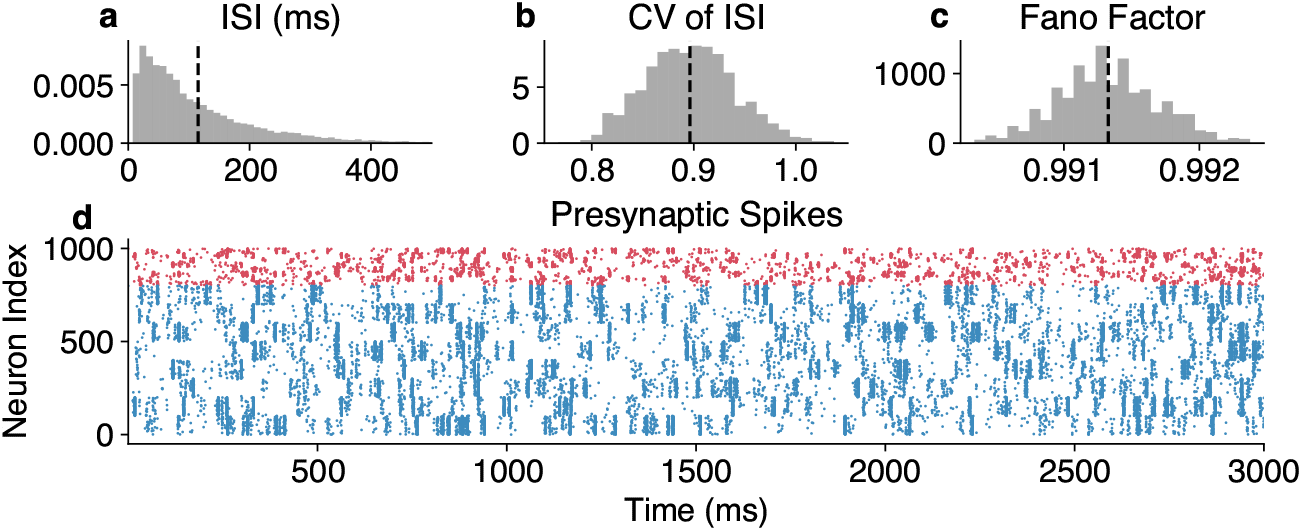
The firing of a feedforward network: **a**. The inter-spike interval distribution. **b**. The distribution of the Coefficient of variation ((CVs) of the ISIs (**c**)The distribution of the Fano Factor, is very close to 1, indicatinng the Poisson input the neurons receive. (**d**) A raster plot visualizing 3 seconds of the network’s activity (Here blue indicates excitatory and red inhibitory neurons)

We then examine a case which is particularly detrimental to the emergence of co-tuning. We introduce unstructured (no assemblies) recurrent connectivity (*p* = 0.5 and *W* = 2) and high noise (noise intensity is set to 0.6).

The resulting distribution of the interspike intervals becomes somewhat skewed, as does the distribution of the CV of the ISI, while the Fano Factor remains very close to 1 (Fig. 8a-c). The network develops occasional large bursts of synchronized activity involving neurons from all groups (Fig. 8d).

**FIG. 8.**
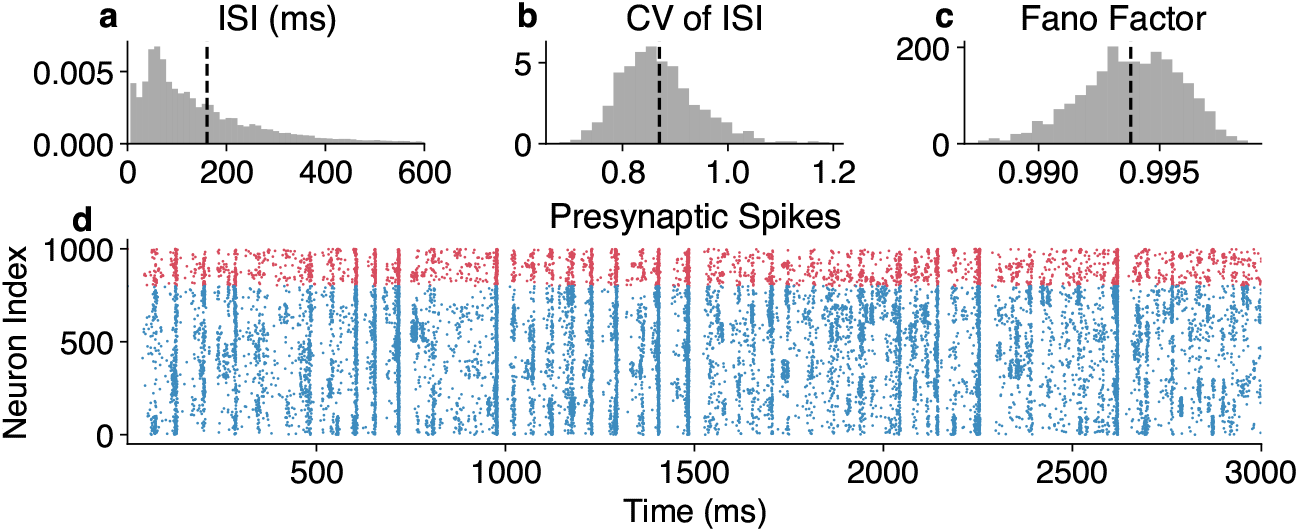
The firing of a network with strong noise (noise intensity = 0.6) and strong random recurrence (*p* = 0.5 and *W* = 2): **a**. The inter-spike interval distribution. **b**. The distribution of the Coefficient of variation ((CVs) of the ISIs (**c**)The distribution of the Fano Factor, remains very close to 1. (**d**) A raster plot visualizing 3 seconds of the network’s activity (Here blue indicates excitatory and red inhibitory neurons)

Finally, we look at the previous network, albeit with the introduction of optimal assembly structure (*r*_*EE*_ = *r*_*EI*_ = 1.0 and *r*_*IE*_ = 1.0 and r_IE_ *r*_*II*_ = 0.4).

This network is characterized by short synchronized firing periods of each group, during which neurons of other groups are largely silent (Fig. 9d). This leads to a bimodal distribution of the mean ISI and a very skewed distribution of the CV of the ISI, which indicates relatively irregular firing (largely due to the combination of bursts and longer periods of silence). Finally, the Fano Factor remains close to 1 (Fig. 9a-c).

**FIG. 9.**
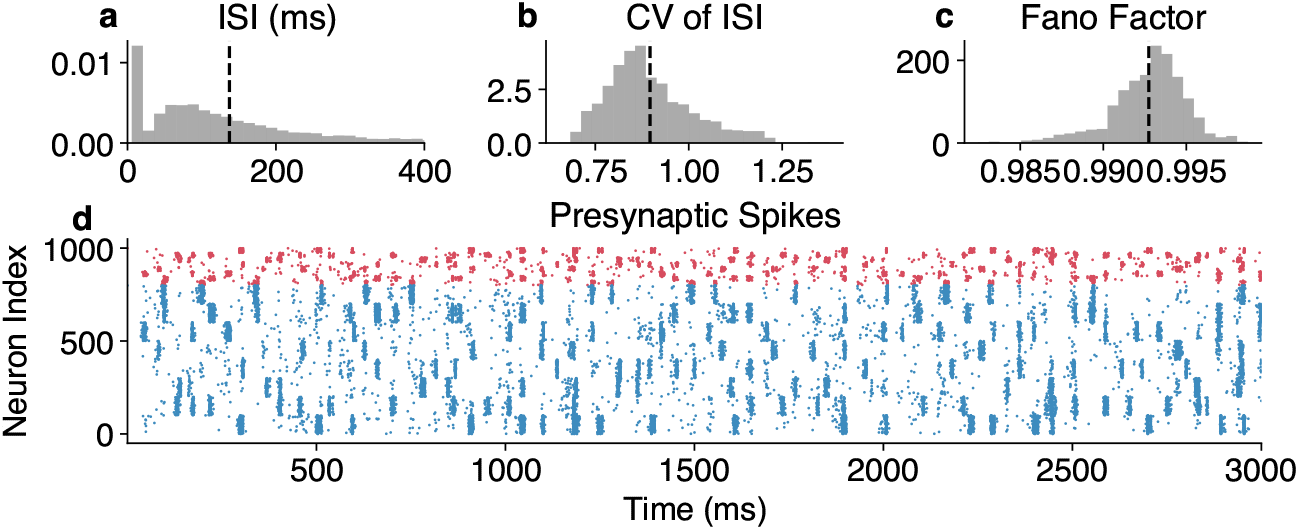
The firing of a network with near-optimal clustering (*r*_EE_ = *r*_EI_ = 1.0 and *r*_IE_ = *r*_II_ = 0.4): **a**. The inter-spike interval distribution. **b**. The distribution of the Coefficient of variation ((CVs) of the ISIs. The average increases slightly compared to the previous networks. (**c**)The distribution of the Fano Factor, remains very close to 1. (**d**) A raster plot visualizing 3 seconds of the network’s activity (Here blue indicates excitatory and red inhibitory neurons)

## Appendix 4 Learned connectivity and the dynamics of the post-synaptic neuron

Since synaptic plasticity depends not only on presynaptic network statistics but also on postsynaptic neuron activity, we examine the learned connectivity of different networks (the ones we examined in the last section) and visualize the activity of the postsynaptic neuron.

At first, the feedforward, low-noise network leads to a very strongly co-tuned E/I connectivity (Fig. 10a) and diverse weights between different groups. The incoming currents are highly correlated (Fig. 10b) and the post-synaptic neuron fires (Fig. 10c) relatively sparsely (⟨ISI⟩ = 0.398, CV_ISI_ = 1.15, Fano Factor = 0.991).

**FIG. 10.**
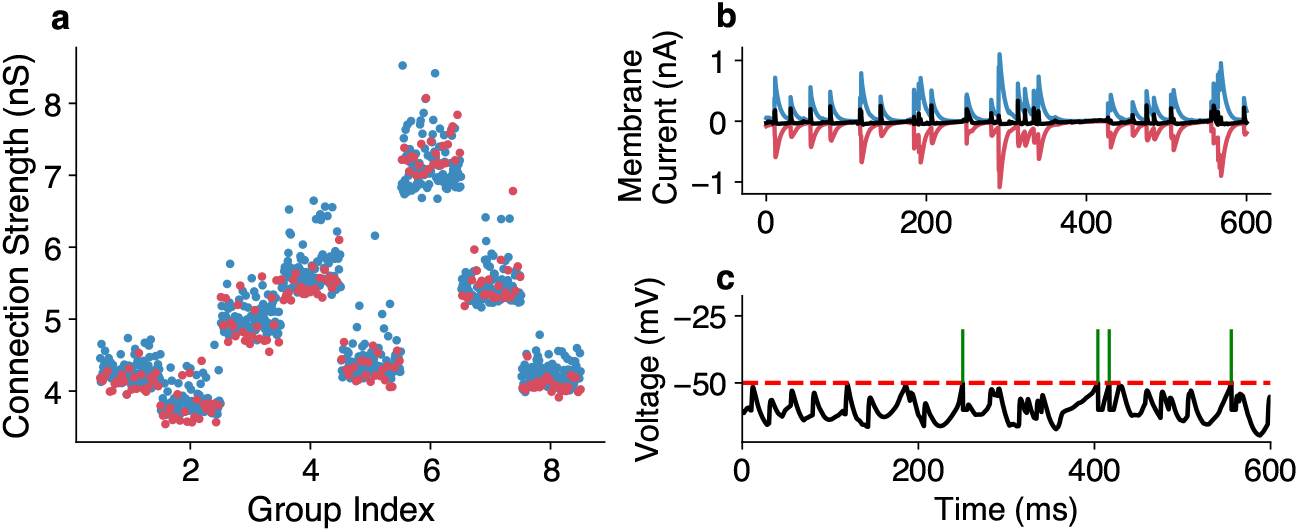
Connectivity and firing of a postsynaptic neuron receiving input from a feedforward network: **a**. The learned connectivity is co-tuned and diverse. **b**. The incoming currents are tightly balanced. **c** The voltage trace and spikes of the readout neuron.

On the contrary, the noisy and recurrent network fails to develop any structure in the learned connectivity. The coordinated firing across groups, visualized in (Fig. 11a), leads to occasional large current influxes (Fig. 11b), but otherwise, the neuron remains relatively tightly balanced. The neuron’s firing (Fig. 11c) becomes slightly more frequent (⟨ISI⟩ = 0.21, CV_ISI_ = 1.23, Fano Factor = 0.991).

**FIG. 11.**
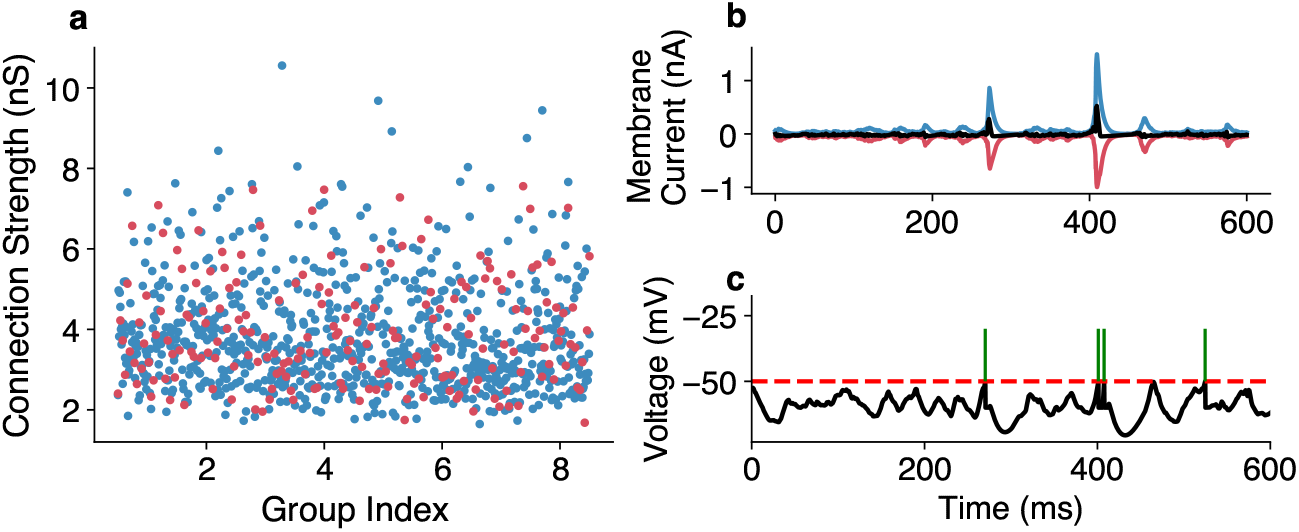
Connectivity and firing of a postsynaptic neuron receiving input from a noisy, recurrent network: **a**. The learned connectivity is completely unstructured. **b**. The incoming currents are tightly balanced despite occasional large influxes. **c** The voltage trace and spikes of the readout neuron.

Finally, the network with the optimized assembly structure, due to the restored statistics of the input (Fig. 12a), develops co-tuned E/I connectivity and relatively diverse weights between groups. The currents arriving to the postynaptic neurons are tightly balanced (Fig. 12b) and the firing of the postynaptic neuron (Fig. 12c) becomes again more sparse and relatively irregular (⟨ISI⟩ = 0.61, CV_ISI_ = 1.14, Fano Factor = 0.993).

**FIG. 12.**
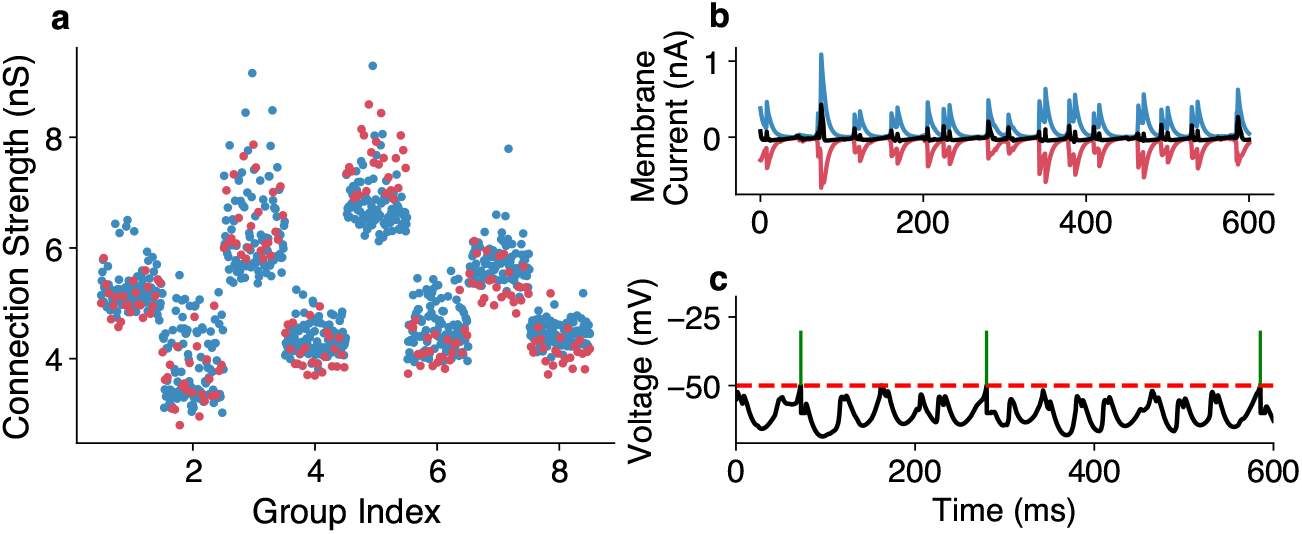
Connectivity and firing of a postsynaptic neuron receiving input from a network with optimal assembly structure: **a**. The learned connectivity is restored to be co-tuned and diverse. **b**. The incoming currents are tightly balanced. **c** The voltage trace and spikes of the readout neuron.

## Appendix 5 The effects of inhomogeneous connectivity are independent of the plasticity protocols’ details

We examined whether our results are dependent on the particular plasticity protocol we used, and we verified that they hold for alternative plasticity mechanisms. Specifically, we tested whether a variety of different plasticity protocols produces co-tuning in the simple feedforward case, whether the effects of noise and recurrence on the resulting connectivity are consistent and whether the inferred optimal fixed pre-synaptic recurrent connectivity restores the ability of the plasticity to produce co-tuning.

Starting with the plasticity protocol we used in our original experiments, we visualize the development of excitatory (Fig. 13a) and inhibitory weights (Fig. 13b), the matching resulting connectivity (Fig. 13c) and the incoming E/I currents to the post-synaptic neuron after convergence (Fig. 13d) in a setting with optimal connectivity for comparison with the other plasticity protocols simulated with the same connectivity and noise levels.

**FIG. 13.**
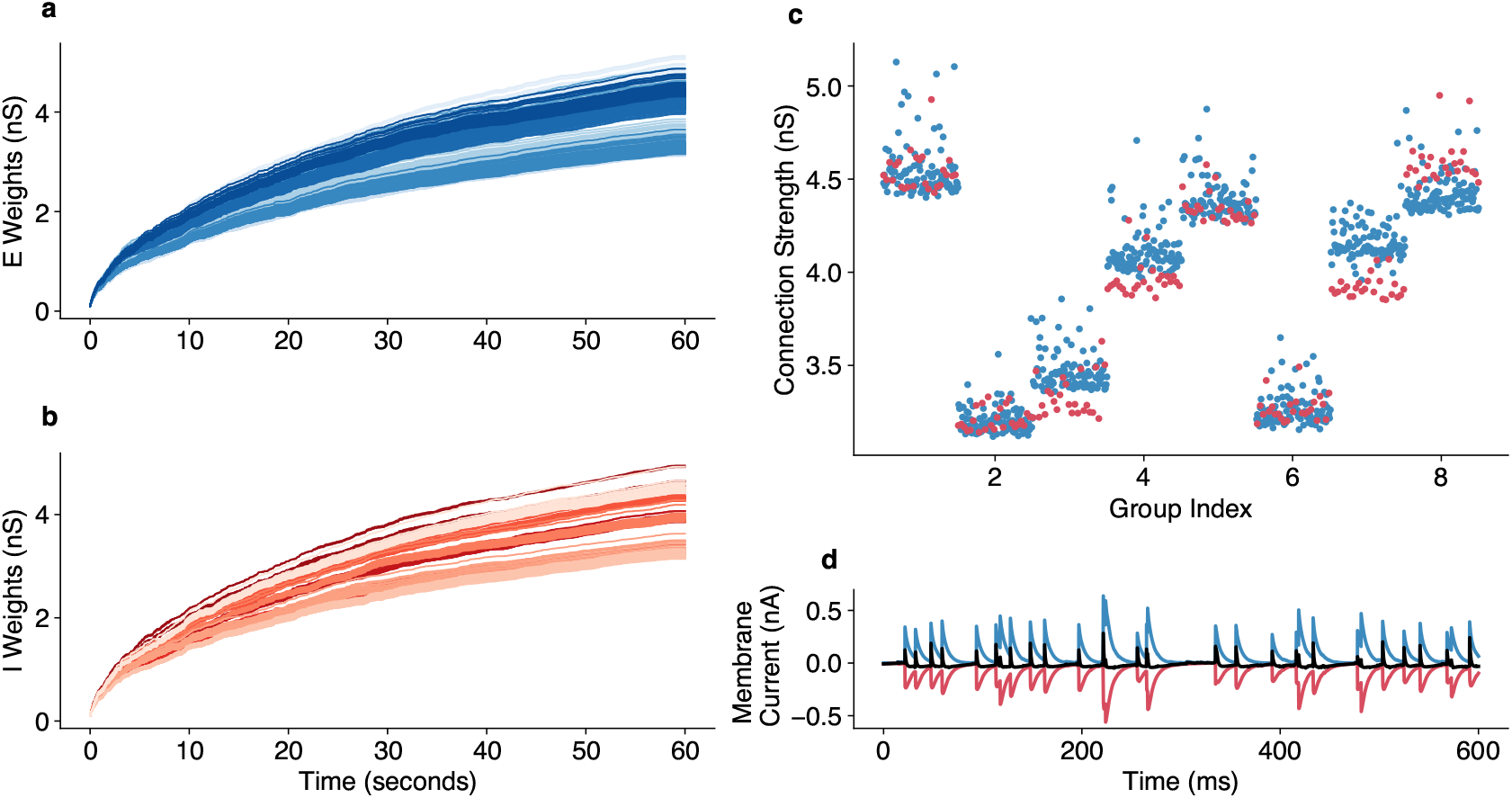
Weight development under the original protocol. The development of **a**. excitatory and **b**. inhibitory weights over a minute of simulation time. **c**. The convergence point of excitatory (blue) and inhibitory weights (red). The inhibitory weights are scaled to account for the slower synaptic constant of the inhibitory synapses and the smaller number of inhibitory neurons. **d**. The resulting E/I currents on the postsynaptic neuron are tightly balanced.

### 1. Excitatory plasticity: Parameters of the Triplet rule

In our experiments presented in the main text, we have used a simplified form of the triplet STDP rule for the excitatory synapses, which relies on identical slow and fast timescales for the pre and postsynaptic traces. We first tested how robust our results are to changes on the LTD/LTP ratio (varying the *A*_*LTD*_ ∈ [0.05, 1.2] and *A*_*LTP*_ ∈ [0.05, 1.0]) as well as the timescale of the fast and slow traces. We found that qualitatively our results are robust to these changes in the plasticity protocol.

Moreover, in order to ensure that our results did not depend on the particular form of the rule we used, we replicated our main simulations using the original implementation of the triplet rule from [48], which implemented different timescales for the pre and postsynaptic fast and slow traces.

Specifically, in this implementation, the pre and post-synaptic activity is tracked by the traces:

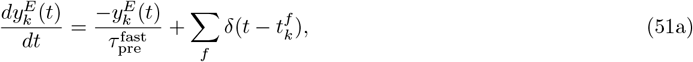

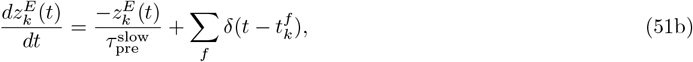

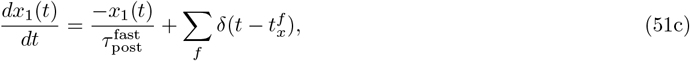

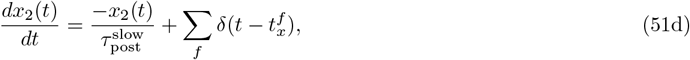

where 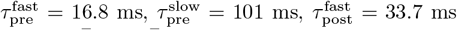 and 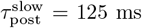 following the parameters from [23, 48]. Additionally, 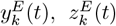 and 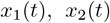 represent the slow and fast traces of the *k*-th excitatory pre-synaptic and the single post-synaptic neuron respectively while 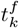 and 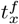are their respective firing times. The function *δ*(*x*) represents a Dirac’s delta. The connection weights are updated upon pre and post-synaptic spiking according to

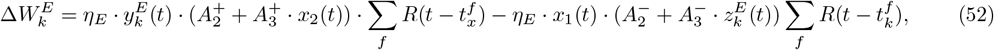

where 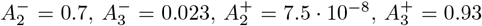 and *η*_*E*_ = 10^−2^, following the parameters of [23, 48]. We see that under this protocol (Fig. 14a, b), the weights develop very similarly to the one we used for our experiments (Fig. 13a, b) under optimal connectivity. Moreover, we see that strong co-tuning emerges (Fig. 14c) as well as tightly balanced E/I currents to the postsynaptic neuron as expected (Fig. 14d).

**FIG. 14.**
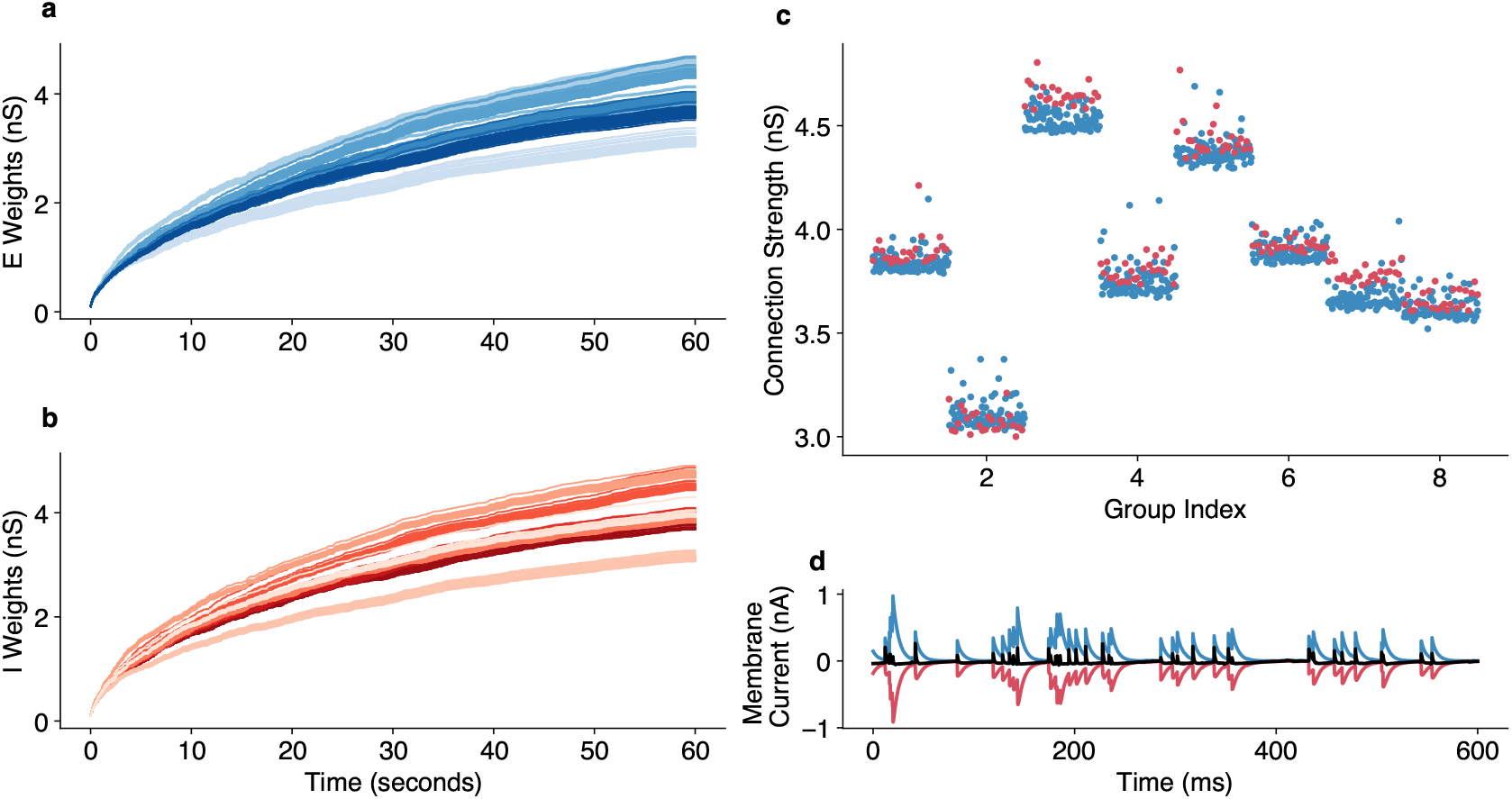
Weight development with the original triplet rule parameters. The development of **a**. excitatory and **b**. inhibitory weights over a minute of simulation time. **c**. The convergence point of excitatory (blue) and inhibitory weights (red). The inhibitory weights are scaled to account for the slower synaptic constant of the inhibitory synapses and the smaller number of inhibitory neurons. **d**. The resulting E/I currents on the postsynaptic neuron are tightly balanced.

### 2. Excitatory plasticity: Triplet vs Pair rule

We further examined whether a different Hebbian learning rule in the excitatory synapses has any impact on our results. In particular, we replaced the triplet rule [48] with a classic spike pair Hebbian rule [84], which relies on a single pre and postsynaptic trace:

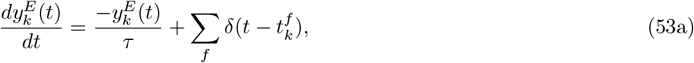

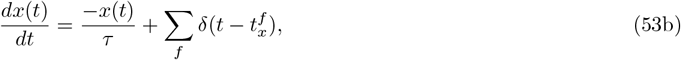

where 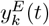 the trace of the *k*-th presynaptic excitatory neuron, *x*(*t*) the trace of the postsynaptic neuron and *τ* = 10 ms the time constant of the decay. The weight update happens as:

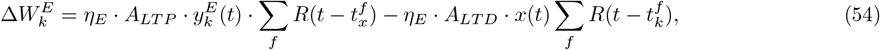

where *A*_*LT P*_ = 1.0 and *A*_*LTD*_ = 0.2 same as for the simplified triplet rule. Using the same competitive, synapse-type specific normalization, we found that in this setting, the weight development as well as the impact of noise and recurrence are similar to our original setting.

### 3. Inhibitory learning rule target rate

We additionally examined several different target rates *ρ*_0_ (ranging from 0 to 6 Hz) for the inhibitory plasticity rule (setting the rate to *ρ*_0_ = 0, i.e., making the inhibitory plasticity into a pure symmetric Hebbian rule, as was used in the original study of the normalization mechanism by [31]). Besides an expected change in the postsynaptic firing rate after the convergence of the weights, we did not observe any other noticeable changes in our findings regarding the emergence of co-tuning and input selectivity.

### 4. Alternative implementations of the competitive Normalization

#### a. Regular vs event-based normalization steps

For reasons of numerical simulation speed, we implemented the weight normalisation in an asynchronous manner. Specifically, we apply a normalisation step after every weight update occurs (i.e., after each postsynaptic spike all connections get normalised and after each presynaptic spike the corresponding connection gets normalised). Since the presynaptic population has a homogeneous firing rate (i.e., all neurons spike approximately the same number of times) all connections are normalised approximately equally often, which makes this implementation behave similarly to implementing the normalisation step on each time step of the simulation.

To test that this assumption is correct, we simulated a network with a regular normalization update, where all connections are normalized simultaneously every time step (*dt* = 0.1 ms). Besides a modification of the rate of the normalization *η*_*N*_ (necessary to counter the higher frequency of normalization in the regular case), we maintain the exact same parameters as in the original experiment.

We observe that in this setting, the weight development happens very similar to the original experiment (Fig. 15a, b), producing co-tuning in cases with optimal input connectivity (Fig. 15c) and tight E/I current balance (Fig. 15d). In summary, we find that the two normalization methods behave similarly for our setting, suggesting that our findings are independent of how the weight normalization is implemented. However, in settings different from ours, where the pre-synaptic rates are largely inhomogeneous, we would expect that the two normalization methods would produce different results (since some weights would be normalized more often than others), which might lead to different weight dynamics. Thus, the numerical convenience of the asynchronous normalization updates cannot be generally used in simulations of other types of networks with potentially very different distributions of firing rates.

**FIG. 15.**
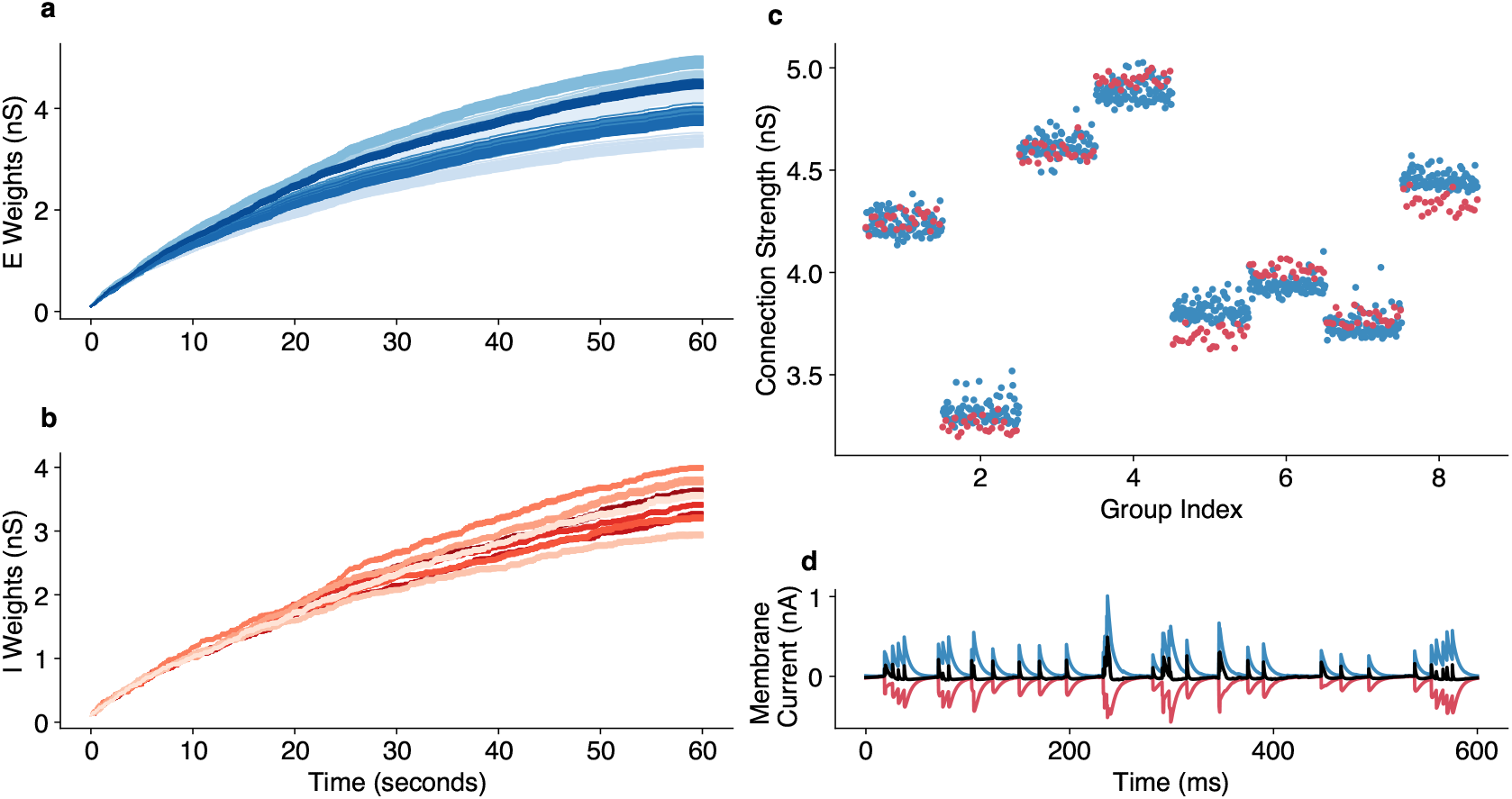
Weight development with a regular normalization update. The development of **a**. excitatory and **b**. inhibitory weights over a minute of simulation time. **c**. The convergence point of excitatory (blue) and inhibitory weights (red). The inhibitory weights are scaled to account for the slower synaptic constant of the inhibitory synapses and the smaller number of inhibitory neurons. **d**. The resulting E/I currents on the postsynaptic neuron are tightly balanced.

#### b. “Soft” vs “Strict” normalization: The impact of the normalization rate

As described in the “Methods” section of the main text, we implement our normalization step as:

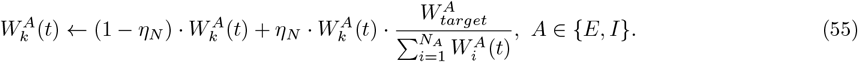

where *η*_*N*_ = 3 10^−3^ to match the equivalent excitatory and inhibitory learning rates. This essentially implements a “soft” normalization, where the total sum of the incoming weights to the postsynaptic neuron is not kept constant in each timestep, but rather pushed towards maintaining a sum as close as possible to the target weight sum over time.

However, if we treated the normalization strictly as the preservation of synaptic resources over time following [31], we would need to set *η*_*N*_ = 1, in order to preserve the exact weight sum constant for each time step. This approach would impose a very strict condition which in combination with the small learning rates of the E and I plasticity, would prevent weight diversification.

One way of countering this is by massively increasing the E and I learning rates, but this tends to make the learning dynamics unstable. Another way of solving this problem is by implementing the normalisation regularly but not on every time step (a solution that has been used previously in [23]), for example, implementing it every 20 ms. This does lead to stable learning, but it also occasionally promotes winner-take-all connectivity, which is not in itself problematic, but may be undesirable for some types of coding. The emergence of winner-take-all connectivity can, in turn, be potentially countered by faster inhibitory plasticity relative to the excitatory plasticity, but finding the exact parameters can involve a fair amount of fine-tuning. For our experiment, the “soft” which can be biologically justified as a competition for synaptic resources among multiple neurons, happening on a very slow timescale, clearly leads to more plausible and stable results.

### 5. Subtractive Normalization with modified input

Finally, in order to fully demonstrate that our results are independent of the exact normalization protocol, we studied both the triplet [48] and pair [84] excitatory Hebbian rules combined with a subtractive normalization mechanism that has been previously used in plasticity studies [23, 33]. Following our setting for the multiplicative normalization, we applied this rule also as a “soft” normalization in the excitatory synapses (the inhibitory synapses are not normalized in this setting):

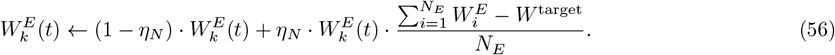

In 2016 [12], it was demonstrated that subtractive normalization only on the excitatory connections will lead to all the weights converging on the same point due to the inhibitory plasticity creating a moving threshold. In order to prevent this collapse of the receptive field, enforced inhomogeneity on the firing rates of different groups is needed. We solved this problem by giving the network’s input as pulses of 100 mS during which some of the input groups firing rate quadruples. This enforces inhomogeneous firing rates, which result in the emergence of stable, diverse, and co-tuned feedforward connectivity (Fig. 16a - c). We also verified that the post-synaptic neuron is tightly balanced (Fig. 16d) and maintains a stable firing rate.

**FIG. 16.**
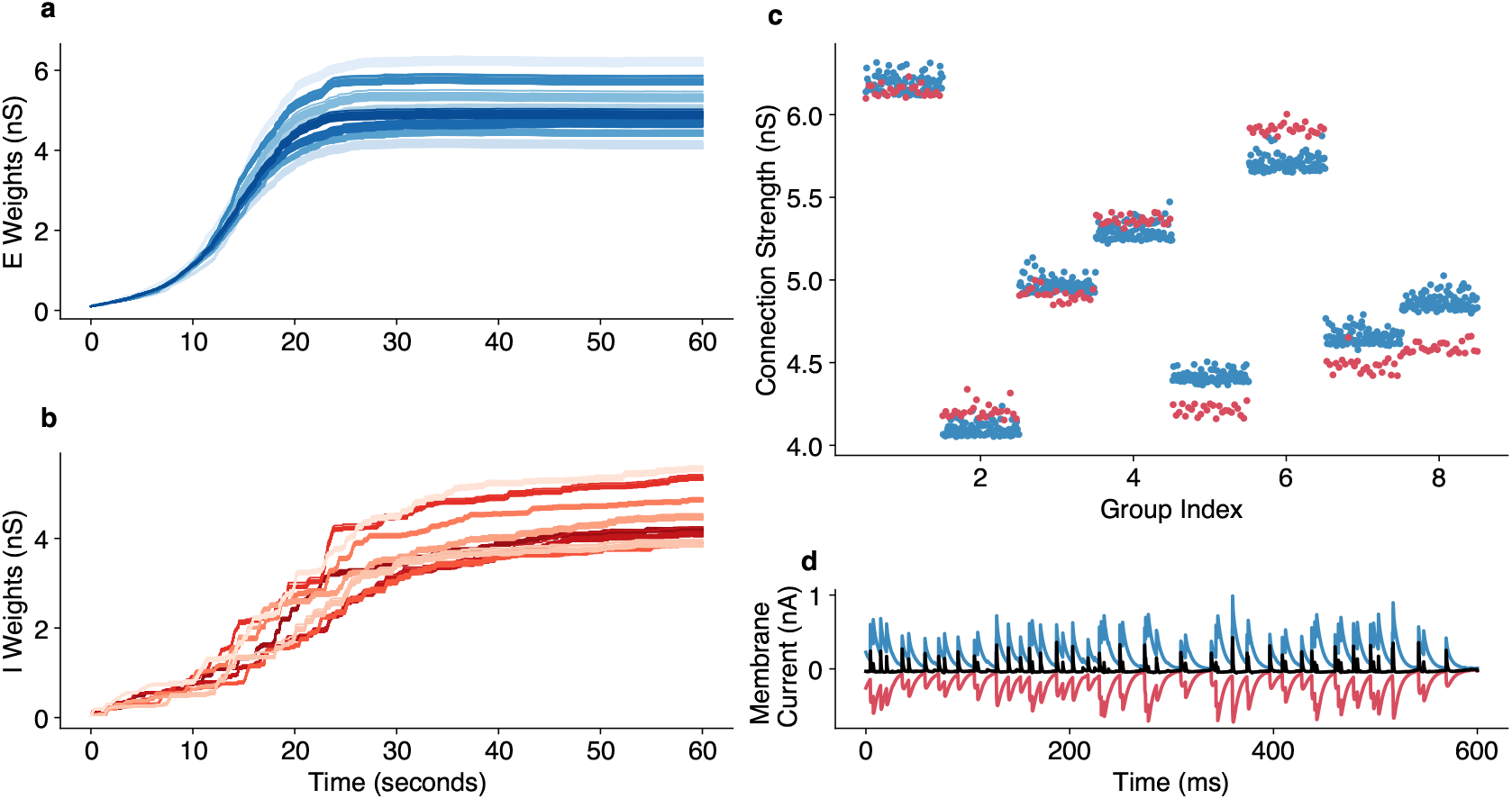
Alternative normalization protocol. The development of **a**. excitatory and **b**. inhibitory weights over a minute of simulation time. **c**. The convergence point of excitatory (blue) and inhibitory weights (red). The inhibitory weights are scaled to account for the slower synaptic constant of the inhibitory synapses and the smaller number of inhibitory neurons. **d**. The resulting E/I currents on the postsynaptic neuron are tightly balanced.

We verified that the co-tuning achieved in this setting, similar to the mechanism presented in the main text, suffers from the introduction of noise and recurrent connectivity. Furthermore, the assembling principles that we derived for the original network seem to have a similarly beneficial effect on this setting, restoring the original covariance structure of the network’s activity and leading to detailed co-tuning between the excitatory and inhibitory feedforward connections.

## Appendix 6 Convergence of weights to an eigenvector of a modified covariance matrix under plasticity

The study that introduced the competitive synapse type-specific normalization [31] analytically predicted that for Hebbian E and I feedforward plasticity, the convergence point of the weights is an eigenvector of the modified covariance matrix (Fig. 17a):

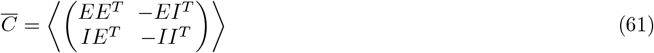

where *E, I* are the activities of the excitatory and inhibitory populations, respectively.

**FIG. 17.**
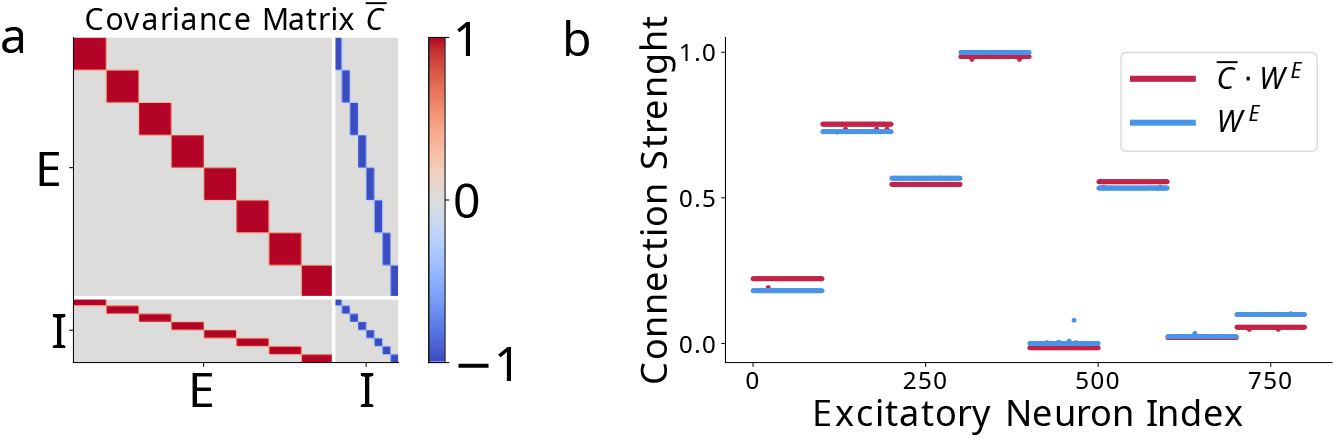
Weights converge to an eigenvector of the covariance matrix. **a**: The estimated covariance matrix for a feedforward network. **b**. We verify that the convergence point of the weights is an eigenvector of the covariance matrix.

We test whether in our modified plasticity protocol (simplified Triplet STDP in E connections, slower normalization), the convergence point can be approximated by an eigenvector of the above matrix.

Specifically, we multiply the converged weight vector with a numerical calculation of the covariance (estimated via binning of the spike trains with a bin size of 1 mS), for different noise and recurrence settings, and we verify that the resulting product is approximately equal to a multiple of the original weight matrix (Fig. 17b). This indicates that despite the differences in the learning protocol, the plasticity converge point is largely controlled by the covariance structure of the population activity.

## Appendix 7 Perturbation of the optimal assembly strengths leads to diverse effects on the network’s activity

To investigate the relative importance of the different connection types, we perturb various assemblies away from the optimal solutions inferred with ABC and check the resulting changes on the network activity as well as how fast the weight co-tuning and diversity deteriorate.

We find that *E* ← *I* assemblies have a strong impact on the resulting network dynamics and, consequently, the statistics that determine the emergence of co-tuning (Fig. 18e). Thus, when *E* ← *I* assemblies are too strong, the network exhibits synchronous behaviour that does not allow discrimination between neuron groups and, consequently, the emergence of co-tuned connectivity. In contrast, when *E* ← *I* assemblies are too weak, one group of neurons is constantly active. This eventually leads to perfect discrimination of only a single input.

**FIG. 18.**
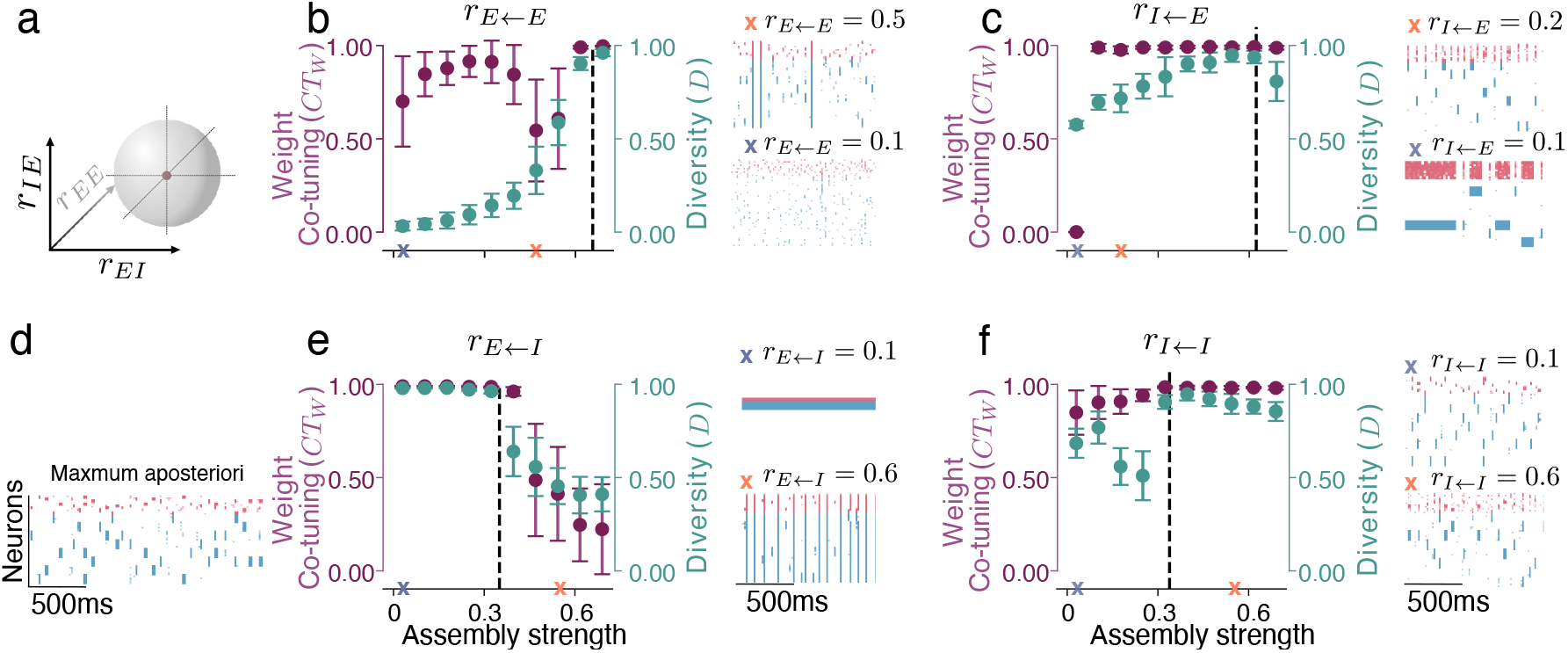
Changes of the various assemblies strengths differently affect weight co-tuning and diversity. **a**. We sequentially vary the value of each assembly strength while keeping the rest of the parameters fixed at the maximum a posteriori (MAP) solution (**d**). **b**. A decrease in the *E* ← *E*, assembly strength (*R*_E ← E_) introduces synchronous burst-like events that jeopardize co-tuning and weight diversity. Further reduction of the *E* ← *E* assembly strength results in sparse, asynchronous spiking, which significantly reduces tuning quality. **c**. Reduction of *I* ← *E* assembly strength first leads to synchronous inhibitory firing across groups, and further reduction leads to persistent activity of the whole inhibitory population combined with bursts of excitatory activity that prevent the development of diversity. **e**. Decreasing the *E* ← *I* assembly strength leads to persistent activation of a single group of neurons (which affects the tuning only marginally). While the increase of the *E* ← *I* assembly strength leads to synchronized behavior of the whole network. **f**. Weakening the *I* ← *I* assemblies decreases the weight diversity by introducing occasional synchronous bursts in the network while strengthening them leads to asynchronous inhibitory activity.

*E* ← *E* assemblies also have a strong effect on the resulting learned connectivity. When they are reduced, the weight diversity rapidly reduces, while the weight co-tuning metric first decreases and then restores. This behaviour is related to two different bifurcations in the population dynamics. First, synchronous full-network bursts emerge on top of the stimulus-induced firing (Fig. 18a). Then, as *E* ← *E* are reduced further, the network enters an asynchronous irregular state (Fig. 18a).

Perturbations of *I* ← *I* and *I* ← *E* assemblies have a weaker effect on the weight co-tuning and diversity. Specifically, when *I* ← *I* assemblies are weakened, the network displays synchronous bursts on top of the stimulus-driven activity that only minimally reduces the weight co-tuning and diversity (Fig. 18d). When *I* ← *E* assemblies are reduced, the weight diversity slowly decreases due to a decrease in the diversity of inhibitory weights. Thus, the activity of the inhibitory population becomes more discoordinated (Fig. 18c). On the other hand, a network with uniform *I* ← *E* connections (i.e., without *I* ← *E* assemblies), shows a dramatic reduction in the weight co-tuning metric because of very strong activation of a single neuron group accompanied by the simultaneous activity of all inhibitory neurons (Fig. 18c).

## Appendix 8 Reduced model calculations

### 1. Derivation of the equations

In this Appendix, we start from the main text Eqs. 14,

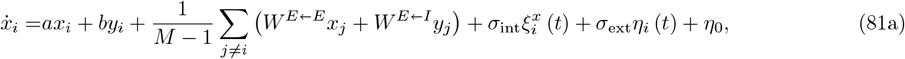

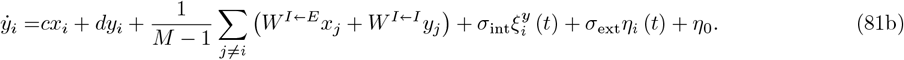

where we consider *M* groups composed by excitatory *x*_*i*_(*t*) and inhibitory *y*_*i*_(*t*) populations (*i* = 1, …, *n*) coupled linearly (see also Methods). Internal noise of each population is represented by 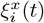 and 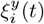 for excitatory and inhibitory populations, respectively. On the other hand, external noise *η*_*i*_(*t*) is shared among both populations. All the noises have zero mean and correlations

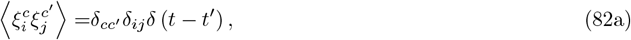

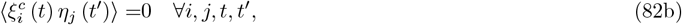

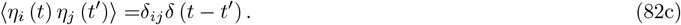

We will derive closed expressions for the correlation coefficients. We would like to remark that ⟨·⟩ means an ensemble average over noise realizations. All stochastic equations are to be interpreted in the Itô convention[85].

First of all, we redefine the noise terms, which will prove convenient later to simplify the algebra. For this reason, we define

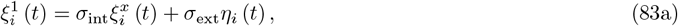

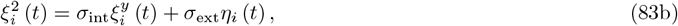

which are Gaussian white noises with zero mean and correlation matrix

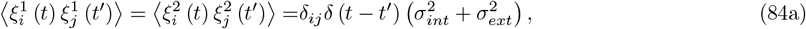

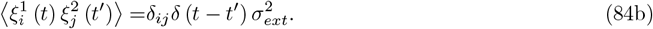

To start with, one can obtain the average values for the stationary rates by applying averages to both sides of equations (81) and imposing the stationary state condition, 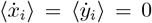. Once this is done, it is immediate to solve the resulting linear system and check that 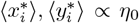, where the star (*) indicates that these values correspond to the stationary state. Hence, making *η*_0_ = 0 the mean values vanish. One can demonstrate that correlations do not depend on *η*_0_, and hence we can make *η*_0_ = 0 without loss of generality. Conceptually, this means just shifting up or down the baseline of fluctuations of the firing rate, which does not affect the fluctuations themselves.

To compute correlations, we need to evaluate the second-order moments between different populations as *x*_*i*_*y*_*j*_ or 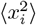. A possible way of doing this is starting from the analytical solution of the multidimensional Orstein-Uhlenbeck process [86]. However, this approach will yield a linear system with *N* (*N* + 1)*/*2 variables to solve for, which are all the elements of the (symmetric) correlation matrix. But all the groups are identical (or *indistinguishable*), so we expect correlations not to depend on the particular population chosen. Therefore, all the equations will be reduced to just 6 covariances: 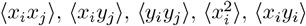 and 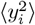.

In this context, it is conceptually simpler to obtain equations for the evolution of the second moments and then evaluate them in the stationary state. Here, we report in detail the computation of two of these moments as an example, giving just the final answer for the other four, which is performed in an analogous way.

First, we define *X*_*ij*_ = *x*_*i*_*x*_*j*_, and then look for the time evolution of *X*_*ij*_, i.e., 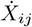. Notice that this is a non-linear change of variables, and thus Itô’s lemma is required. The lemma tells us that if we have a change of variables *z* = *z*(*x*), then one has to include the second-order terms in the expansion,

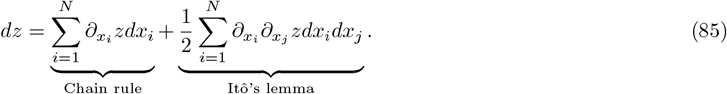

The terms *dx*_*i*_ can be obtained as 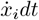. It is important to remark that in this procedure noise terms are rewritten as the differential of the Wiener processes, i.e., *η*_*i*_(*t*)*dt* = *dW*_*i*_. After applying the Itô lemma above, only terms up to order *dt* should be taken into account. Notice that 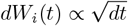 [86]. Finally, one just divides again by *dt* to recover the stochastic differential equation and applies the ensemble average.

For *X*_*ij*_, this reads as

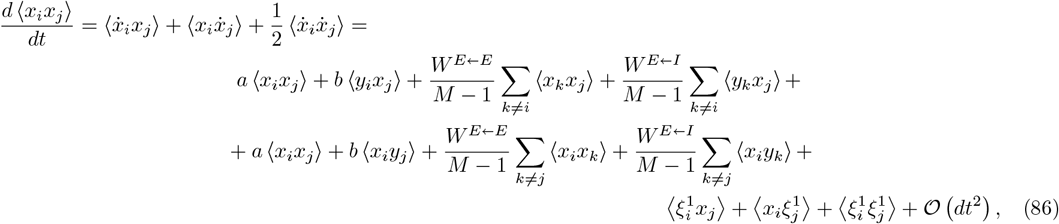

where all the averages between the noise and the variable yield 0, due to Itô’s prescription. The next step is to simplify the sums involving correlations. As discussed above, since clusters are indistinguishable, all the terms are exactly the same. However, the dummy index *k* will also take the value of the fixed index, *k* = *j*, and this has to be taken into account separately since the in-group is different to the between-group one. Then,

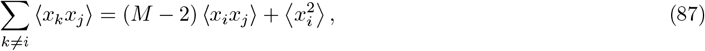

allowing us to simplify the equation. At this step we simplify the notation by letting *X*_*ij*_ = ⟨*x*_*i*_*x*_*j*_⟩, *Z*_*ij*_ = ⟨*x*_*i*_*y*_*j*_⟩, 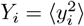, etc., leading to

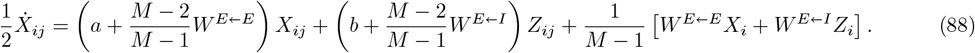

The same procedure can be repeated for all the other correlations, such as

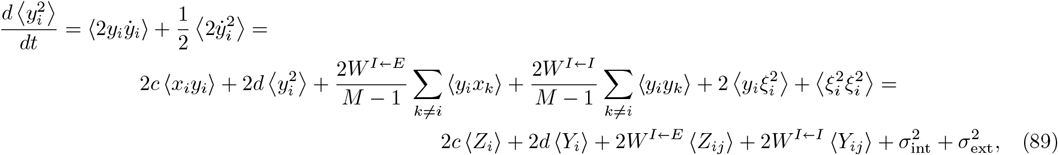

where now the correlation between noises yields a non-vanishing value. This operation is repeated with all the remaining terms, in order to find a linear system of differential equations with 6 variables and 6 equations,

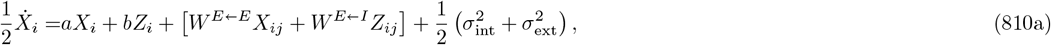

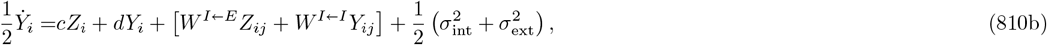

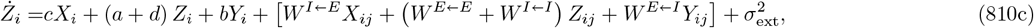

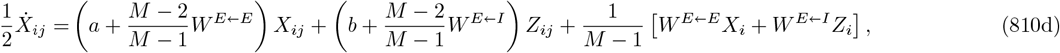

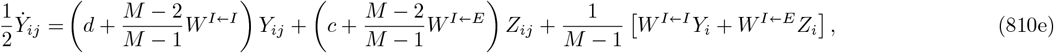

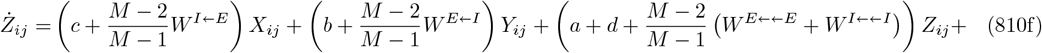

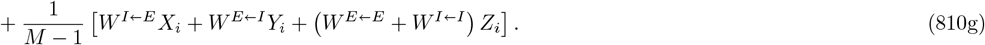

This system can be solved in the stationary limit when all the derivatives of the left-hand side are zero. From these, one is able to obtain the Pearson correlation coefficients. Correlation with itself is always unity, thus there are only four coefficients remaining: the correlation between excitation and inhibition inside a group 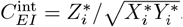, and all three between-group correlations, 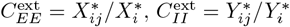, and 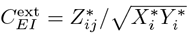.

### 2. Solutions for the homogeneous network

In some special cases, it is possible to give a simple solution in closed form for the correlation coefficients. One example is the homogeneous network: when all weights are identical, and an intrinsic decay is added to both the excitatory and inhibitory populations (i.e., with *c* = *W* ^*E* ← *E*^ = *W* ^*I* ← *E*^ = +*W*, *b* = *W* ^*I* ← *E*^ = *W* ^*I* ← *I*^ = −*W* and *a* = *W* − 1, *d* = −*W* − 1) the solution reads

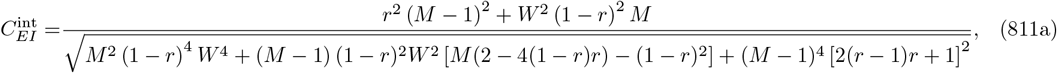

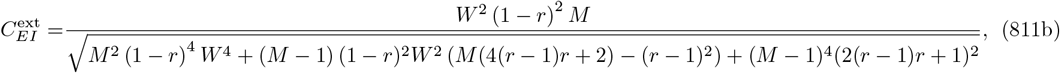

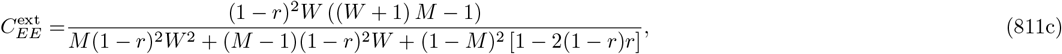

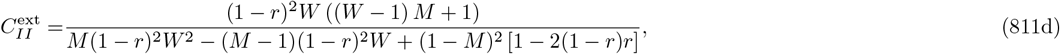

where we defined *r* as the signal-to-noise ratio, *σ*_int_ = *rσ, σ*_ext_ = (1 − *r*)*σ*. This analytical solution has some interesting features. First, notice it does not depend on the total amount of noise *σ* that the system receives, but only on the ratio between external and internal noise. Second, if *W*→∞ all correlations go to 1, making the diversity between groups vanish. Expanding in series around *ϵ* = 1*/W* = 0, one can see that all coefficients are *C* = 1 − 𝒪 (1*/W* ^2^) for large coupling.

It is also possible to study the limiting values of the noise. *r* = 1 makes all the between-group correlations equal to zero, while coupling determines the in-group value. On the other hand, when *r* ≪ 1, one gets

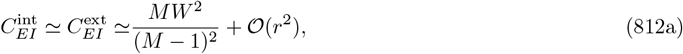

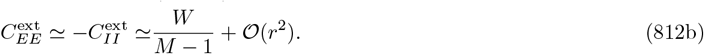

meaning that the external correlations grow linearly with the coupling, but quadratically with the signal-to-noise ratio: a small increase in coupling needs to be followed by a larger increase in signal intensity in order to recover the previous tuning. As a result, the coupling has a larger impact on tuning than the signal-to-noise ratio, an effect that can be measured in the full spiking network.

Finally, we see that between-group correlations also tend to zero as the limit *M* → + ∞ is taken, since in that case, the input that a module receives from all others becomes just white noise. A finite number of clusters (or finite connectivity among them) is thus required for tuning.

## Appendix 9 Clustering optimisation

Optimization of clustering for a fully connected network can be done by minimizing a loss function that depends on the correlations. A simple possibility is to employ minimum squares,

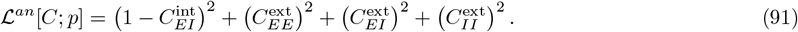

The solution and associated optimal correlations are shown in 19. There are several key remarks following from this figure:

1. Even for very large clustering and extremely low signal-to-noise ratio, the clustering is able to provide an in-group correlation close to unity combined with low between-group correlations, thus ensuring co-tuning.
2. Inhibition to excitation never clusters. Inhibitory neurons act over excitatory individuals regardless of their cluster.
3. Excitatory connections are clustered. In particular, excitatory connections always project to inhibitory neurons in their own cluster but not to other ones. Excitatory-to-excitatory connectivity is also strongly clustered, except for large coupling.
4. Inhibition controls excitation for large *W*. If one keeps highly clustered excitation and increases the coupling, the dynamics of single modules become unstable at a critical value *W*_*c*_(*r*). However, the network can remain stable if the excitatory clustering is reduced and the amount of inhibition in the group increases, which can be accomplished by increasing *r*_*II*_.
5. When the signal-to-noise ratio is close to one, clustering becomes mostly irrelevant, since the system is driven by the external input, which allows co-tuning easily.

Notice that the optimization algorithm automatically finds solutions where the equations are well-defined –i.e., where the system reaches a stationary state– thus selecting to increase the inhibitory clustering when *W* goes over the instability threshold (Fig. 19).

**FIG. 19.**
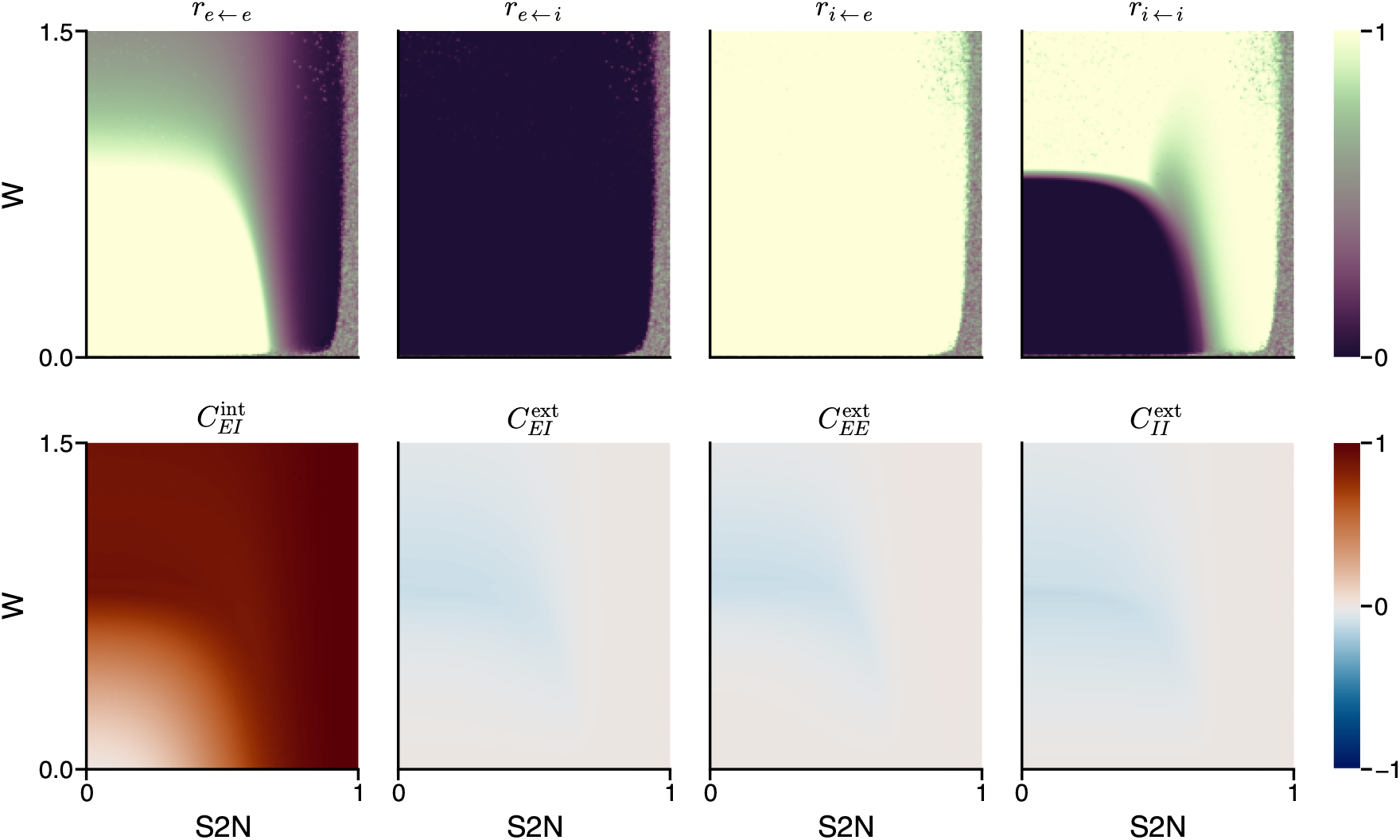
Optimical clustering for the analytical results. The selected values of the clustering lead to large decorrelations between different groups. Large correlations in-group can be achieved for a large set of recurrence and external noise. In this case, highly clustered excitation and homogeneous inhibition are recovered.

Therefore, the analytical approach is able to find a good candidate for optimal clustering depending on the network dynamics. Although it cannot be directly applied to the spiking network, which is able to display richer dynamics, it tells us that, as a rule of thumb, excitatory clustering should be as high as possible while avoiding crossing an instability threshold. If this happens, inhibition needs to be increased.

## Appendix 10 The inferred connectivity structure encourages competition between assemblies

The distribution of optimal assembly strengths we identify (Fig. 20a), consists in very strong excitatory assemblies and much weaker inhibitory assemblies, leading to strong excitatory connections among neurons of the same input group and more spread-out inhibitory connections that also target neurons from other groups.

**FIG. 20.**
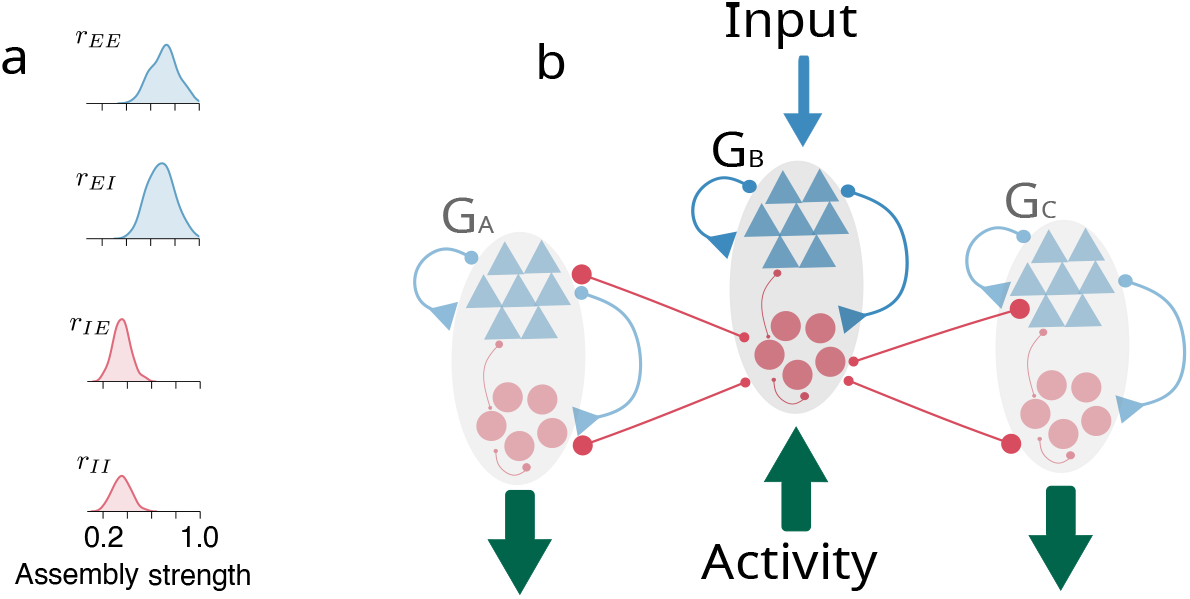
**a** The posterior distrubution of assembly strengths. It promotes very localized excitatory and relatively broad inhibitory connectivity. **b** A diagrammatic representation of how the inferred connectivity is tuned for decorrelating the activities of different groups

In our network, this global inhibition setting promotes competition between different assemblies which generates the desired correlation structure in the population activity. Essentially, when a group receives an external input, the strong excitatory connections within the group (projecting to both E and I in-group neurons) will lead to high activity of all the neurons (both E and I) in the group. However, unlike the E connections, which largely target other neurons inside the group, the I connections (targeting both E and I neurons) are mostly directed toward the neurons of other groups, which means that the high inhibitory activity inside the group would lead to the suppression of the activity (in both E and I neurons) of other groups.

## Appendix 11 Tables of parameters

We mostly used the neuron model parameters from the original inhibitory STDP paper [26].

| Network Model |  |  |
| --- | --- | --- |
| Symbol | Description | Value |
| $N$ | Number of neurons | 1000 |
| $N_E$ | Number of E neurons | 800 |
| $N_I$ | Number of I neurons | 200 |
| $M$ | Number of input groups | 8 |
| $g_{\text{leak}}$ | Leak conductance | 10 nS |
| $V_{\text{rest}}$ | Resting potential | -60 mV |
| $V_{\text{reset}}$ | Reset potential | -60 mV |
| $V_{\text{th}}$ | Spiking threshold | -50 mV |
| $V_E$ | Excitatory reversal potential | 0 mV |
| $V_I$ | Inhibitory reversal potential | -80 mV |
| $C_m$ | Membrane capacitance | 200 pF |
| $\tau_{\text{ref}}$ | Absolute refractory period | 5 ms |
| $\tau_E$ | Decay time constant of E conductance | 5 ms |
| $\tau_I$ | Decay time constant of I conductance | 10 ms |
| $\overline{g_E}$ | E weight scaling constant | 1.4 nS |
| $\overline{g_I}$ | I weight scaling constant | 3.5 nS |

| Plasticity Rules |  |  |
| --- | --- | --- |
| Symbol | Description | Value |
| $\tau_1^{\text{estdp}}$ | Slow eSTDP timescale | 50 ms |
| $\tau_2^{\text{estdp}}$ | Fast eSTDP timescale | 10 ms |
| $\eta_E$ | eSTDP learning rate | 0.0025 |
| $A_{\text{LTP}}$ | Long term potentiation amplitude | 1.0 |
| $A_{\text{LTD}}$ | Long term depression amplitude | 0.2 |
| $\tau^{\text{istdp}}$ | iSTDP timescale | 10 ms |
| $\eta_I$ | iSTDP learning rate | 0.01 |
| $\rho_0$ | iSTDP target firing rate | 3 Hz |
| $\eta_N$ | Normalization learning rate | 0.003 |
| $W_{\text{target}}^E$ | Excitatory normalization target | 5.0 |
| $W_{\text{target}}^I$ | Inhibitory normalization target | 5.0 |

| ABC Optimization |  |  |
| --- | --- | --- |
| Symbol | Description | Value |
| $\alpha$ | weight of in-group correlation | 0.1 |
| $\beta$ | weight of between-group correlations | 0.3 |

## Notes

### Competing Interest Statement

The authors have declared no competing interest.

### Summary of Updates

We have added a new section on the impact of structure in a fully-plastic network. All figures are revised for clarity.

